# Parallel monitoring of mRNA abundance, localisation and compactness with correlative single molecule FISH on LR White embedded samples

**DOI:** 10.1101/2020.07.14.203562

**Authors:** Susanne Kramer, Elisabeth Meyer-Natus, Hanna Thoma, Achim Schnaufer, Markus Engstler

## Abstract

Single mRNA molecules are frequently detected by single molecule fluorescence in situ hybridisation (smFISH) using branched DNA technology. While providing strong and background-reduced signals, the method is inefficient in detecting mRNAs within dense structures, in monitoring mRNA compactness and in quantifying abundant mRNAs.

To overcome these limitations, we have hybridised slices of high pressure frozen, LR White embedded cells (LR White smFISH). mRNA detection is physically restricted to the surface of the resin. This enables single molecule detection of RNAs with accuracy comparable to RNA sequencing, irrespective of their abundance, while at the same time providing spatial information on RNA localisation that can be complemented with immunofluorescence and electron microscopy, as well as electron tomography. Moreover, LR White embedding restricts the number of available probe pair recognition sites for each mRNA to a small subset. As a consequence, differences in signal intensities between RNA populations reflect differences in RNA tertiary structures, and we show that the method can be employed to probe for mRNA compactness. We apply LR White smFISH to answer some outstanding questions related to trans-splicing, RNA granules and mitochondrial RNA editing, using trypanosomes and their versatile RNA biology as a model system.

## INTRODUCTION

Accurate and quantitative determination of RNA localisation is essential to understand RNA function. Often, it is important to determine the percentage of a given RNA species in a certain sub-compartment of a cell. The study of nuclear export for example requires accurate monitoring of RNA distribution between nucleus and cytoplasm. Another example is the study of nuclear or cytoplasmic RNA granules, phase-separated aggregates of RNA and protein with still poorly understood functions.

<u>F</u>luorescence <u>i</u>n <u>s</u>itu <u>h</u>ybridisation (FISH) visualises abundant RNA by hybridisation with a fluorescently labelled DNA oligo. This method is only semi-quantitative, as hybridisation efficiency is highly sensitive to minor changes in conditions and also differs between subcellular environments. Higher accuracy is provided by single molecule FISH (smFISH). Each RNA molecule is quantified as one dot, regardless of its brightness, and differences in hybridisation efficiency are thus compensated for. smFISH is done by one of two methods. In standard smFISH, the RNA molecule is probed with a set of DNA oligos coupled to fluorophores. In branched DNA technology, the RNA molecule is detected by a set of up to 20 probe pairs, each consisting of two adjacent oligos that further hybridise in several steps with other oligos to generate a large branched structure of DNA finally linked to fluorophores ^1^. Branched DNA technology has two major advantages in comparison to standard smFISH: (1) background fluorescence is massively reduced, as each signal depends on hybridisation of two oligos next to each other (binding of just one oligo will not result in a signal) and (2) the amplification of the signal results in labelling with up to 400 fluorophores per probe pair and thus reduces the minimal size of the target sequence to about 50 nucleotides (recognition site of one probe pair). This allows probing for very small RNAs or distinguishing RNAs with minor sequence differences, for example alternative splicing products.

The disadvantage of branched DNA technology is its massive requirement in space to build the DNA branch. Each mRNA is labelled with up to 8,000 fluorophores (400 fluorophores label each of the up to 20 probe pairs). The exact nature of the fluorophores is proprietary, but, for example, 8,000 Alexa Fluor™ 488 fluorophores would correspond to a combined mass of >5,000 kDa, not including the DNA part. In particular mRNAs within protein-dense structures are therefore difficult to detect. The Affymetrix technology (Thermo Fisher Scientific) uses a protease to circumvent the problem, on the cost of loss of most cellular structures, such as antigens needed for immunofluorescence. While the method appears to work relatively well in large, cultured mammalian cells, its limits become in particular apparent when working with smaller cells such as yeast and many protozoans, that have a cell wall or dense cytoskeleton underneath the plasma membrane. In trypanosomes for example, there is a huge variance in fluorescence intensities between mRNA molecules within the same cell ^2,3^ and the same is observed on recent smFISH images from *Plasmodium* ^4^. Almost no yeast publications exist that use branched DNA technology.

Here, we set out to establish a method that allows quantitative studies of RNAs without loss of cell integrity, regardless of cell architecture, shape and origin. Instead of chemical fixation and protease digestion, high-pressure-frozen cells are embedded into LR White (=LR White smFISH). The solid resin reduces probing physically to the surface of the microtome slice, and this has several advantages: (1) the signal becomes independent of its sub-cellular environment and thus allows probing within protein-dense structures; (2) the amount of RNA molecules accessible to the probes is greatly reduced, allowing the detection of even highly abundant RNAs as single molecules; (3) structures and dimensions of the cells are preserved, avoiding artefacts caused by fixation and protease digestions; and (4) images can be correlated with electron microscopy.

As a model system, we used the flagellated protozoan *Trypanosoma brucei*, the causative agent of African Sleeping sickness and one of the causes of the related cattle disease Nagana. The parasite has several advantages for establishing LR White smFISH. (1) With ~2×20 μm the trypanosome cell is large enough to allow RNA localisation while still being small enough to image an entire cell. (2) The cell is highly polar and detailed spatial information is provided by the specific positions of several single-copy organelles. For example, trypanosomes have one single mitochondrion and its mitochondrial DNA (the kinetoplast) can be stained with DAPI and localises to a very precise position close to the basal body. This unique cell architecture facilitates intracellular localisation studies of RNA. (3) Two different trypanosome life cycle stages, the bloodstream form (BSF; mammalian stage) and the procyclic form (PCF; tsetse fly stage) can be cultured that differ in morphology and metabolism and in their major cell surface proteins. Here, these differences serve as essential controls for the specificity of the RNA smFISH. For example, BSF cells express the variant surface glycoprotein (VSG) as their major surface protein, while PCF cells express the procyclin EP1 and variants of it. mRNA levels of the mRNA encoding the respective cell surface protein are tightly regulated and only detectable in the respective life cycle stage. (4) Trypanosomes have a fascinating mRNA metabolism with many open questions that can be addressed by LR White smFISH. Some features of mRNA metabolism in trypanosomes are highly conserved with other eukaryotes, for example, the parasites possess a range of RNA granules (dynamic, dense structures of RNA and protein that are phase-separated from the rest of the cytoplasm ^5^) that are very similar to the ones defined in yeast and human ^6–8^. RNA content and proteome of trypanosome starvation stress granules has been defined by granule purification ^6^. Other features of trypanosome RNA biology are rather unique to trypanosomes ^9^: for example, dozens to hundreds of functionally unrelated mRNAs are polycistronically transcribed from epigenetically defined promoter regions ^10^ and subsequently processed by trans-splicing the capped, 39 nucleotide long miniexon of the spliced leader RNA (SL RNA) to each 5’ end; this is coupled with polyadenylation of the 3’ end of the upstream transcript ^11–13^. As such polycistronic transcription prevents high transcription rates; highly abundant mRNAs are either encoded by multiple gene copies (*TUBULIN*, ribosomal protein mRNAs) ^14,15^ or are transcribed by RNA Polymerase I (*VSG, EP1*) ^16^. As a consequence of the unregulated polycistronic transcription, gene expression is regulated almost entirely posttranscriptionally. Trypanosomes are also famous for a particular type of RNA editing: most mitochondrially encoded mRNAs are changed by a complex RNA editing machinery that adds or removes uridylyls. This process is directed by guide RNA molecules (gRNAs) that are encoded by the massive DNA network of the single mitochondrion (the kinetoplast); in extreme cases more than 50% of the RNA sequence is changed post-transcriptionally ^17^.

In this work, we have established LR White smFISH, using trypanosomes as a model system. We show that the method can accurately quantify and localise RNAs of both low and high abundance, independent of their subcellular localisations. We combine LR White smFISH with both immunofluorescence and electron tomography. Importantly, as only a fraction of each mRNA is exposed to the resin’s surface, signal intensity reflects the mRNA’s 3D structure, and we show that this method can be used as a novel tool to probe for mRNA compactness. We use the method to investigate some outstanding questions in trypanosome RNA biology: we quantify and localise the SL RNA, mRNAs encoding ribosomal proteins as well as edited and unedited mitochondrial mRNAs encoding *A6* and *COXIII*.

## MATERIALS AND METHODS

### Trypanosomes

Monomorphic bloodstream stage parasites of the *Trypanosoma brucei brucei* strain Lister 427 (MITat1.2, expressing VSG 221) were cultured in HMI-9 ^18^ at 37°C and 5% CO_2_. *Trypanosoma brucei brucei* Lister 427 procyclic cells were cultured in SDM-79 ^19^. For starvation (generation of stress granules), PCF cells were washed once in PBS (phosphate buffered saline; 137 mM NaCl, 2.7 mM KCl, 10 mM Na_2_HPO_4_; 1.8 mM KH_2_PO_4_) and incubated for two hours in PBS. For the generation of akinetoplastic cells, BSF trypanosomes expressing an L262P mutant version of the F_1_F_O_-ATPase gamma subunit 20 were treated with 10 nM ethidium bromide for 3 days.

### Affymetrix smFISH on chemically fixed cells

RNA detection in chemically fixed cells was done with the QuantiGene® ViewRNA ISH Cell Assay Kit, Glass Slide Format (Thermo Fisher Scientific), as previously described ^2^.

### Affymetrix smFISH and immunofluorescence on LR White embedded cells

High pressure freezing and LR White embedding of trypanosome cell pellets was done as previously described ^3,21^. 100 nm sections of either PCF or BSF trypanosomes were cut with an ‘ultra Jumbo Diamond Knife’ (Diatome AG) and placed on a poly-lysine slide in the order of cutting, to allow subsequent 3D reconstruction (if required). Affymetrix smFISH was done on these slides essentially following the protocol of the QuantiGene® ViewRNA ISH Cell Assay Kit, Glass Slide Format (Thermo Fisher Scientific), starting with the detergent step, with the following variations: (i) the protease step was omitted (we observed no differences), (ii) we used a self-made washing buffer (0.1xSSC (saline sodium citrate); 0.1% SDS) for all but the last washing step in each of the washings to reduce costs, (iii) the incubation with DAPI at the end was done for 30 minutes, followed by 3 washing steps in PBS. Details on all Affymetrix probe sets are listed in Supplementary Table S1.

For combination with immunofluorescence, the DAPI staining was omitted and slides were kept in PBS overnight. The next day, slides were successively incubated in 20 mM glycine in PBS (15 min), 1% BSA in PBS (15 min), first antibody diluted in 0.1% BSA in PBS (rabbit anti 221 VSG), four times in PBS (4x 5 min), secondary antibody diluted in 0.1% BSA in PBS (Alexa Fluor 488, 594 or 647 goat anti-rabbit, 1:100, 30 min), four times in PBS (4x 5 min), DAPI in PBS (1 μg/ml, 30 min) and four times in PBS (4× 5 min). Samples were embedded in Prolong Diamond antifade (Invitrogen). Note that while the RNA signal was stable for weeks, we observed that the DAPI signal was not stable on LR White embedded cells and imaging is thus best done no later than one day after the experiment.

### Detection of stress granules on LR White embedded samples

We found branched DNA technology (e.g. the Affymetrix poly(A) probes) not suitable for stress granule staining, neither on fixed cells nor on LR White embedded cells. Instead, granules can be stained using fluorescent oligos.

In chemically fixed trypanosomes, starvation stress granules can be detected by visualising the accumulation of mRNAs either by hybridisation with an oligo(dT) or with an oligo antisense to the miniexon sequence, both directly coupled to any fluorophore (this work, ^6,22^). On LR White embedded cells we found oligo(dT) most effective. Importantly, oligo(dT) needs to be coupled to a red fluorophore (Cy3); with a green fluorophore, granules could not be visualised, because the LR White resin has a high auto-fluorescence.

### Light microscopy and image analysis

Imaging of fluorescent samples was mostly done with a DMI6000 wide-field microscope (Leica Microsystems, Mannheim, Germany) and in some cases with a custom build TILL Photonics iMic microscope equipped with a sensicam camera (PCO AG); in both cases we used a 100x oil objective. Stacks with 5-15 images in 100 or 140 nm distances were recorded for LR White FISH and stacks with 40 images a 140 nm were recorded for imaging of chemically fixed cells. Images were deconvolved using Huygens Essential software (SVI, Hilversum, The Netherlands) and are either presented as single plane or as Z-stack projection (sum slices). The VSG protein signal (Figure 6A) was recorded on a Zeiss Elyra S.1 structure illumination microscope with a 63× oil objective and an sCMOS-Camera (PCO Edge 5.5); a single plane image is shown. Image analysis was done using available tools of Fiji ^23^. mRNA dots were identified by the ‘find maxima’ function. Note that samples of LR White embedded trypanosomes have no ‘thickness’ as probing is only on the surface. We still recorded stacks of 5-15 images to collect out-of-focus fluorescence for deconvolution. Note that most images are shown in false colours to optimise visibility for colour-blind readers.

### Scanning electron microscopy and correlation with light microscopy

After completion of light microscopy, samples were prepared for electron microscopy. Cover slips were removed by incubation in PBS for one hour, samples were dried, and slides were cut with a diamond pen to smaller pieces to fit into the scanning electron microscope. For contrasting, samples were incubated in 2% aqueous uranyl acetate for 15 min followed by 3 washing steps in water, incubation in 50% Reynolds’ lead citrate ^24^ in water for 5 min and two washing steps in water. After drying, samples were prepared for SEM and imaged as described ^21^. All images were montaged into an aligned stack of stitched images using the TrakEM2 plugin of Fiji ^23^. Correlation with the fluorescent images was done manually using the free and open source vector graphics editor Inkscape (version 0.91; http://www.inkscape.org), with nuclei as intrinsic landmarks.

## RESULTS

### Overview of RNA targets used in this work

We have selected a range of different trypanosome RNA targets to establish LR White smFISH (Table 1). We include seven cytoplasmic mRNAs with different features: (i) the abundant house-keeping mRNA ***α-TUBULIN***; (ii) the abundant ***RPL7a*** mRNA that is underrepresented in starvation stress granules ^6^; (iii) ***HEL67/DBP1***, an mRNA of average abundance; (iv) the abundant mRNAs encoding the trypanosome cell surface proteins **VSG** (specific for the BSF stage) and **EP1** (specific for the procyclic stage); and (v) the two very large mRNAs ***GB4*** and ***FUTSCH***, that both contain repetitive sequences that allow probing over the entire length. In addition, we included a **poly(A)** probe to detect total mRNAs, and a probe to specifically detect the **SL RNA**, the substrate for the trans-splicing reaction. To study RNA editing, we included probe sets targeting either the edited or unedited version of the mitochondrial localised mRNAs encoding the ATP synthase F_O_ subunit 6 (**A6**) and the cytochrome *c* oxidase subunit III (**COXIII**). Details on the probe sets can be found in Supplementary Table S1.

**Table 1:** Overview of mRNA targets used in this work.

| target | localisation | special feature | specificity controls (this work) | abundance (mRNAs/cell), PCF |  |
| --- | --- | --- | --- | --- | --- |
|  |  |  |  | <sup>25</sup> | this work |
| **all mRNAs** | cytoplasm (nucleus) | poly(A) probe | oligo(dT) competition | 611<br>13 | N.D. |
| *RPL7A* | cytoplasm (nucleus) | underrepresented in stress granules <sup>6</sup> | antisense probe | 274 | 268 |
| *α-TUBULIN* | cytoplasm (nucleus) | abundant house keeping |  | 810 | N.D. |
| *HEL67/DBP1* | cytoplasm (nucleus) | average abundance |  | 21 | 33 |
| *EP1* | cytoplasm (nucleus) | abundant in PCF | life cycle stage specific | 572 | 582 |
| *VSG (MITat1.2)* | cytoplasm (nucleus) | abundant in BSF | life cycle stage specific | N.D. | N.D. |
| *GB4* | cytoplasm (nucleus) | very large mRNA, low abundant |  | 3 | 9.8<sup>*1</sup> |
| *FUTSCH* | cytoplasm (nucleus) | very large mRNA, low abundant |  | 4 | N.D. |
| **A6 edited** | mitochondrion | Pan-edited |  | N.D. | 18<sup>*2</sup> |
| **A6 unedited** | mitochondrion | Pan-edited | ethidium bromide | N.D. | 660<sup>*2</sup> |
| **COXIII edited** | mitochondrion | Pan-edited | ethidium bromide | N.D. | 64<sup>*2</sup> |
| **COXIII unedited** | mitochondrion | Pan-edited | ethidium bromide | N.D. | 1211<sup>*2</sup> |
| **SL RNA** | nucleus | substrate for trans-splicing |  | N.D. | 1100 |

<sup>\*1</sup> includes large fraction of transcription and decay intermediates
<sup>\*2</sup> includes an unknown number of editing intermediates

### Specific detection of single abundant mRNAs

To establish LR White smFISH, a pellet of PCF trypanosome cells was high-pressure frozen, embedded in LR White and cut into 100 nm slices (Figure 1A). The slices were immobilised on poly-lysine slides. Slides with chemically fixed cells served as control (Figure 1B). DAPI stain confirmed that LR White embedded cells have random orientation (the cytoplasm is visible by DAPI background stain in the overexposed image in Figure 1A). Where nuclei coincide with the surface of the cut they are mostly visible as rings surrounding the single nucleolus, indicating that the DAPI stain cannot penetrate into the resin (2D stain) (Figure 1A). In contrast, chemically fixed cells lie flat on the slide, and the nucleolus causes only a slight reduction in DAPI signal, as the DNA underneath and above the nucleolus is still stained (3D stain) (Figure 1B). Kinetoplasts, the DNA of the single mitochondrion, like nuclei are visible in LR White embedded cells only in those cases where they coincide with the surface of the cut.

**Figure 1:**
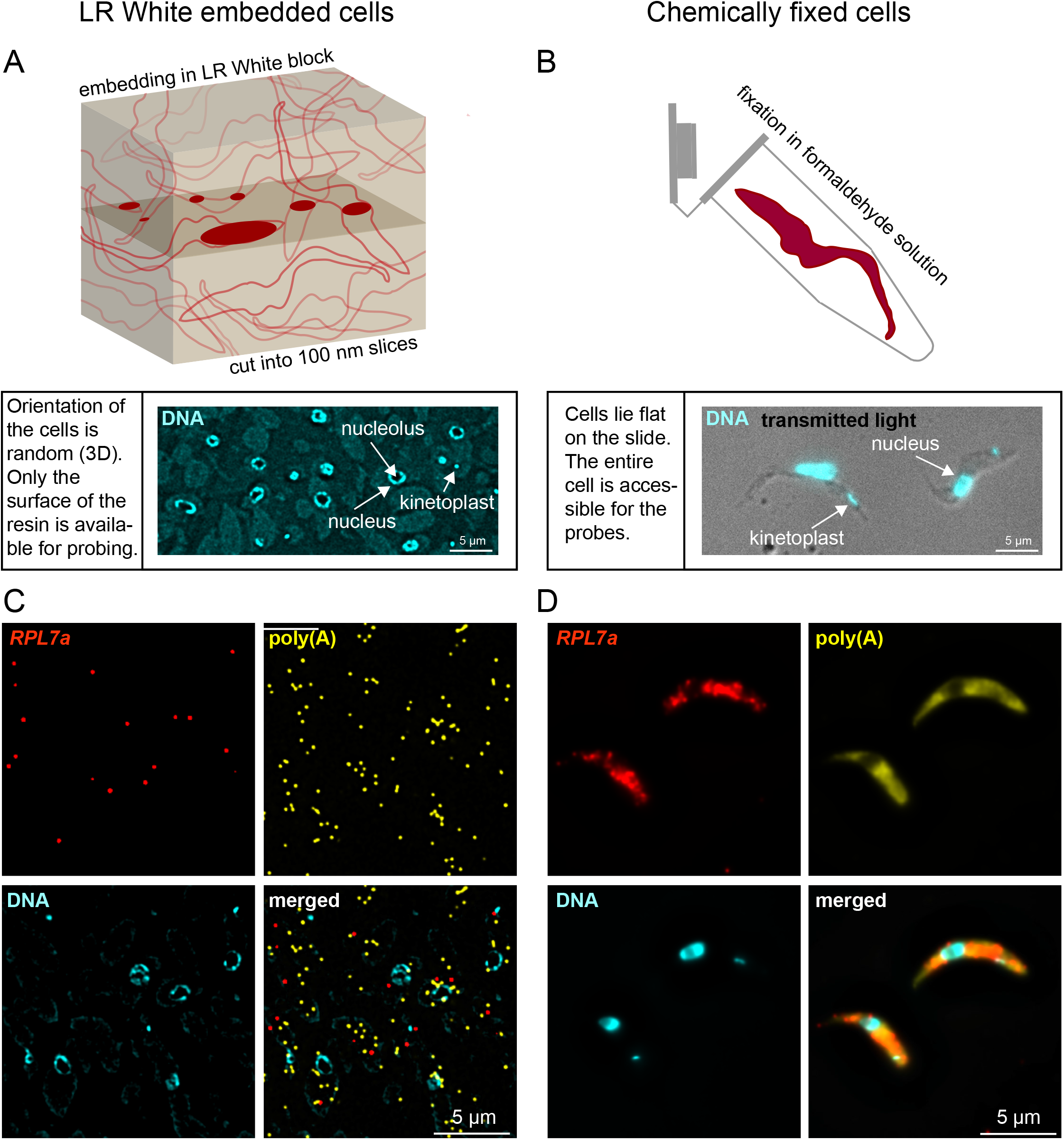
Establishment of LR White smFISH in trypanosomes. **(A)** Top: schematic of a block of LR White with embedded trypanosomes. The plane of cutting is shown, and the surface exposed parts of the cells are shown in solid dark red. Bottom: 100 nm slices of LR White embedded trypanosomes were transferred to a poly-lysine slide. A typical DAPI stain of such a slide is shown slightly overexposed, to visualise the cytoplasm of the cell (background stain) in addition to the nuclei and kinetoplasts. **(B)** Cells were chemically fixed in formaldehyde solution (top, schematic) and transferred to poly-lysine slides. Bottom: an overlay image of transmitted light and DAPI is shown. Slices of LR White embedded PCF trypanosomes **(C)** or chemically fixed PCF trypanosomes **(D)** were probed for *RPL7a* (red) and poly(A) (yellow). DNA was stained with DAPI. Images are shown as deconvolved Z-stack projections.

Slides were incubated with a probe set antisense to the abundant mRNA encoding the ribosomal protein RPL7a (red) and with a probe set antisense to poly(A) (yellow) using the ViewRNA™ ISH Cell Assay Kit (Thermo Fisher Scientific). In LR White embedded cells, individual fluorescent dots were detectable with both probe sets (Figure 1C). In contrast, in chemically fixed cells, individual *RPL7a* mRNA molecules were hardly distinguishable, and the poly(A) signal was equally distributed throughout the cell (Figure 1D).

We performed several control experiments to verify the specificity of the signals. To control for the specificity of the *RPL7a* signal, samples were probed with an *RPL7a* sense probe set. The number of dots detected with the sense probe was with 68 dots / 28,738 poly(A) dots 40 times reduced in comparison to the number of dots detected with the *RPL7a* antisense probe set (2091 dots / 21494 poly(A) dots) (Supplementary Figure S1). To exclude the detection of genomic DNA, we examined the intracellular localisation of all dots originating from the sense *RPL7a* probe (which can detect DNA but not RNA). With 51 cytoplasmic and 16 nuclear dots, we observed no enrichment of dots in the nucleus and thus no evidence for DNA probing. To control for the specificity of the poly(A) signal, we included an oligo(dT) probe in the hybridisation; this oligo massively reduced the poly(A) signal, while the presence of an unrelated oligo had no effect (Supplementary Figure S2). As a further control for the specificity of the method, we probed LR White embedded BSF and PCF trypanosomes for one mRNA that is almost exclusively expressed in the BSF stage (*VSG*) and one that is specific for the PCF stage (*EP1*). While both mRNAs were readily detectable in the expected life cycle stage, the reverse probing gave almost no signal (Supplementary Figure S3).

The data show that abundant mRNAs can be detected as single molecules on LR White embedded samples and that probes are target-specific.

### Quantifying surface-exposed probe pair recognition sites

In chemically fixed samples, each mRNA is detected by the full set of up to 20 probe pairs, with some reductions due to space constraint. Consequently, the detection of an individual mRNA molecule is largely independent of the number of probe pairs used for the probing. In LR White embedded samples, probes can only detect mRNA on the surface of the slice. This causes not only a reduction in the number of detectable mRNA molecules, but also a reduction in the number of probe pairs that can hybridise to each mRNA molecule, as only part of it will be on the surface of the cut. This is demonstrated by the fact that colocalisation between *RPL7a* and poly(A) on LR White embedded slices is a rare event (Figure 1C), even though the majority of *RPL7a* molecules has a poly(A) tail.

To estimate the number of probe pairs that hybridise to an mRNA on LR White embedded samples, we probed the 5’ end of the *RPL7a* mRNA with seven adjacent probe pairs in three different colours: the first probe pair in the first colour (shown in red), the adjacent two probe pairs in the second colour (shown in yellow) and the last four probe pairs in a third colour (shown in cyan) (Figure 2A). The number of mRNA dots increased with the number of probe pairs (Figure 2B). We found 75% of all dots to be of single colour, thus, recognised by 1 (red), 1-2 (yellow) or 1-4 (cyan) probe pairs only. The remaining 25% of the dots were of multiple colours. Only 4% of all dots were of triple colour and thus recognised by four or more probe pairs. About half of the dots recognised by the single red probe pair were not recognised by another probe pair. Taken together, the data show that the number of probe pair recognition sites available for hybridisation with a probe set of seven probe pairs is small, mostly between 1 and 2, rarely extending four. This was further confirmed by staining *RPL7a* simultaneously with identical probe sets of 20 probe pairs in two different colours, shown in red and yellow (Figure 2C): only 46% of mRNA dots were of dual colour. In contrast, in chemically-fixed cells almost all mRNAs detected by such competition staining were dual-coloured, as most of the 20 probe recognition sites are available (Figure 2D and ^2^). These 46% of double coloured dots in the probe competition assay are in best agreement with, on average, four of the twenty recognition sites being accessible to the probes (the theoretical number of double coloured dots for four probe sets is 45.8, Supplementary Table S2). The higher number in accessible probe pair recognition sites in comparison to the triple colour experiments is expected, as 20 probe pairs were used rather than seven: a stretch of 20 probe pair recognition sites is more likely to pass the surface of the resin more than once than a stretch of 7 probe pair recognition sites.

**Figure 2:**
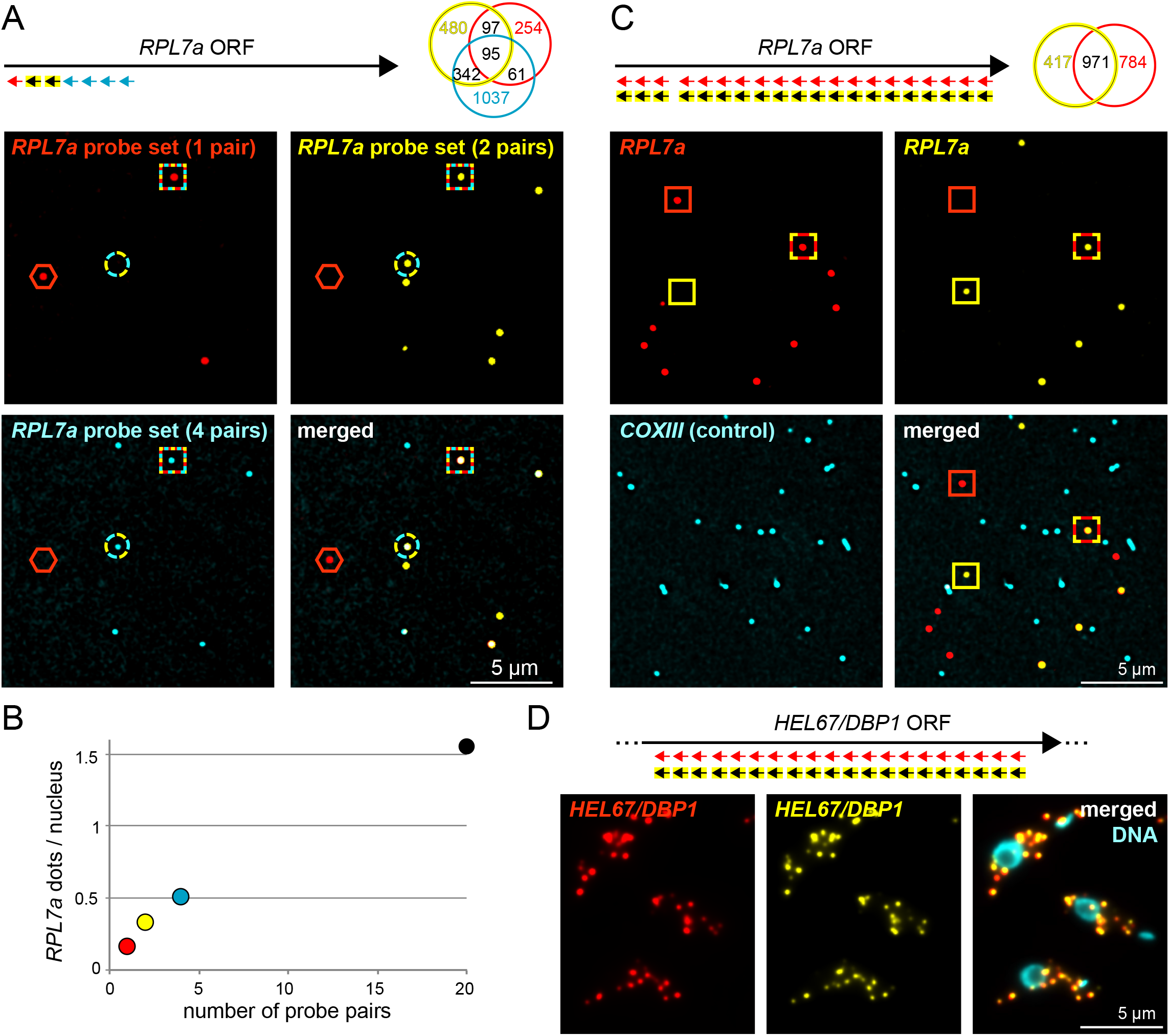
Accessibility of the probes. **(A)** A sample of LR White embedded PCF trypanosomes was hybridised with the tri-colour probe set antisense to *RPL7a*, as indicated. A representative image is shown, with red hexagons, cyan-yellow circles and red-cyan-yellow squares pointing to single, double and triple colour dots, respectively. The numbers of the differently coloured dots are shown as Venn diagram. **(B)** The data shown in Figure 1A (20 probe pairs) and Figure 2A (1, 2 and 4 probe pairs) were used to quantify the number of detectable *RPL7a* molecules, in dependency of the number of probe pairs. The number of visible nuclei was used as a mean of calibration (note that the DAPI stain is not shown in Figure 1A, but was done). The total number of nuclei analysed was 3017 for the sample probed with 1, 2 and 4 probe pairs and 1353 for the sample probed with 20 probe pairs. **(C)** In a competition assay, slices of LR White embedded procyclic trypanosomes were co-probed with 20 probe pairs in two different colours (shown in red and yellow). A representative image is shown, with red, yellow and red-yellow stained dots marked with red, yellow and red-yellow squares, respectively. Quantifications of the differently coloured dots are shown as Venn diagram. With 44.7% double coloured (=red+yellow) dots, these data are in closest agreement with an average of four probe pair recognition sites being exposed for probing (theoretical number = 45.8%, see Supplementary Table S2 for details on this calculation). A probe set antisense to the mitochondrial mRNA *COXIII* served as a negative control (shown in cyan) and showed only 1.6% ‘colocalisation’ with *RPL7a* (4548 cyan dots were analysed). Data of a second experiment were very similar (371 yellow, 650 red and 721 double coloured dots (=41.2%)). We observed a minor bias towards the red signal in both experiments that we cannot explain; there could be minor differences in hybridisation efficiency between the probe sets caused by the different fluorophores. **(D)** Chemically fixed cells were co-probed with 20 probe pairs antisense to *HEL67/DBP1* in two different colours (shown in red and yellow). Representative images are shown. *HEL67/DBP1* was chosen here because of its average abundance; probing for *RPL7a* would not have allowed the detection of individual mRNA molecules (compare Figure 1D).

The data show that the number of probe pair recognition sites affects mRNA detection and needs to be corrected for in any quantification. This is different to the situation in chemically fixed cells. Only a small subset of probe pair recognition sites is exposed to the surface of the resin and thus available for probing.

### Quantifying mRNA levels

To test whether the method allows accurate quantification of mRNAs levels, LR White embedded trypanosomes were probed for five different mRNAs with the aim to compare the results with available data from RNA sequencing ^25^ (Figure 3, Table 1). Next to *RPL7a* (see above), we probed for *EP1* and for *α-TUBULIN*. The *α-TUBULIN* mRNA was chosen for calibration purposes, as it does not contain repetitive sequences and, with on average 810 mRNAs per cell ^25^, is one of the most abundant mRNAs, reducing the error of quantification in both RNA sequencing and smFISH. As representatives of mRNAs of lower abundance we included *HEL76/DBP1* as well as the large mRNA *GB4*, the latter being recognised by a probe set antisense to a highly repetitive sequence spanning 22,389 nucleotides, equalling 468 probe pair recognition sites. All samples were co-probed with an Affymetrix probe antisense to poly(A) and the DNA was stained with DAPI (Figure 3A). For comparison, the same probe sets were used to perform smFISH on chemically fixed cells (Figure 3B).

**Figure 3:**
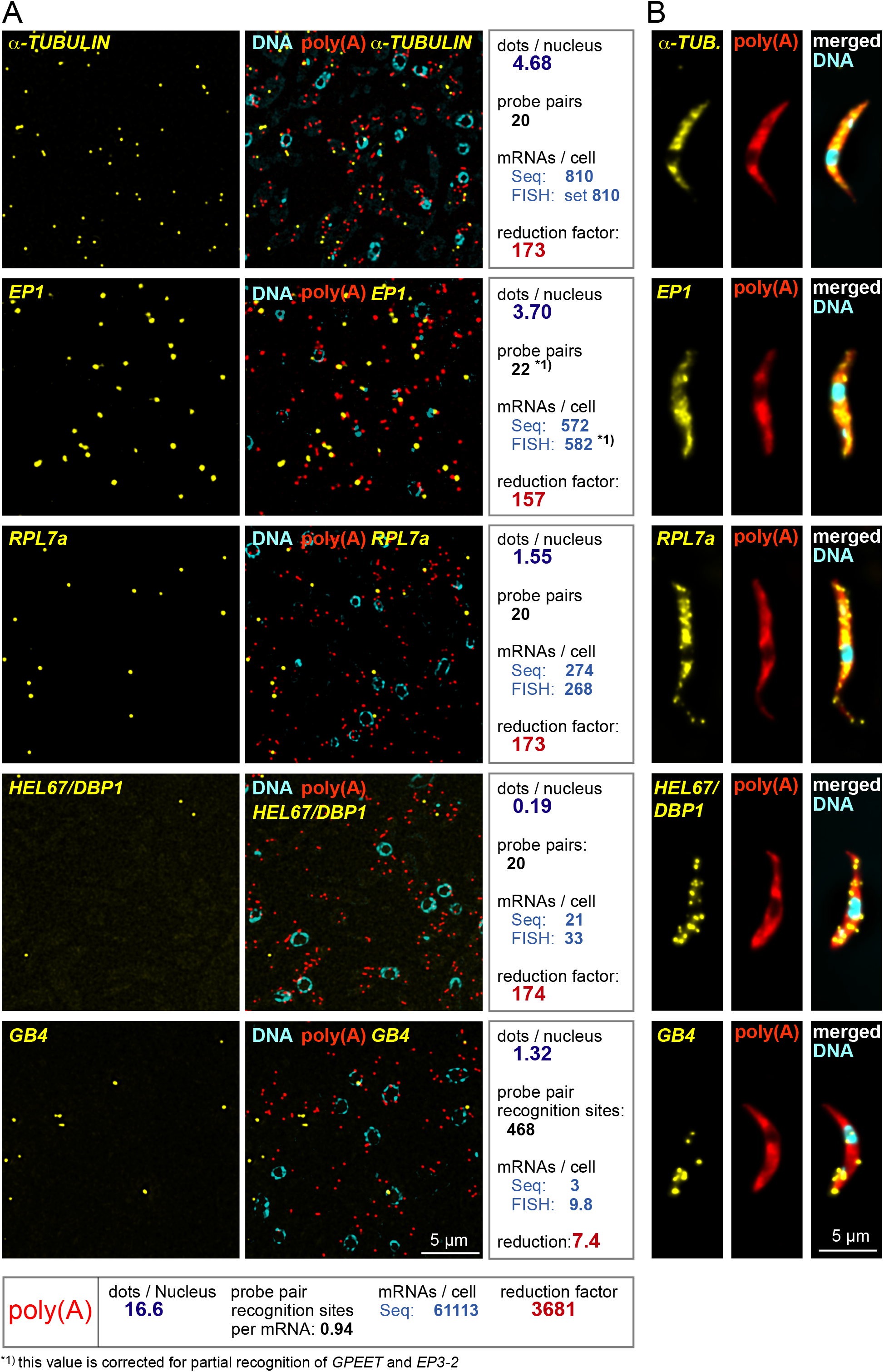
Quantification of mRNAs by LR White smFISH. **(A)** smFISH was performed on LR White embedded samples of PCF trypanosomes using probe sets against the indicated mRNA targets. DNA was stained with DAPI. For analysis, mRNA dots were counted and correlated to the number of nuclei that serve as a proxy for cell number. The total number of mRNA dots used for the analysis was 6082, 4476, 2091, 248, 1792 and 64,058 for *TUBULIN*, *EP1*, *RPL7a*, *HEL67/DBP1, GB4* and poly(A), respectively. Using *TUBULIN* for calibration, the data were compared with RNA sequencing (‘Seq’) data from reference ^25^ to calculate the numbers of mRNAs/cell, with the assumption that an increase/decrease in the number of probe pair recognition sites proportionally changes the number of FISH dots. For *EP1*, one further correction was done: because of sequence similarity in the procyclin gene family, a subset of the 16 *EP1* probe pairs also recognise the related mRNAs *EP2*, *GPEET* and *EP2-3*. Seven probe pairs hybridise to *EP2*, which is of low abundance (7.5 fold less than *EP1* ^25^) and can be neglected, but three probe pairs hybridise to each *EP3-2* and *GPEET*, which are both about as abundant as *EP1* ^25^. Therefore, 22 probe pair recognition sites (16 for EP1, 3 for EP3-2 and 3 for GPEET) were used to calculate *EP1* abundance. The reduction factor is here defined as the quotient of the calculated number of mRNA molecules per cell and the actual number of mRNA dots per nucleus detected by smFISH. Data of one experiment are shown; similar results were obtained in a second experiment with *TUBULIN*=4.88 dots/nucleus, *EP1*=4.12 dots/nucleus, *RPL7a*=1.60 dots/nucleus, *GB4*=1.33 dots/nucleus and *HEL67/DBP1*=0.19 dots/nucleus, analysing data corresponding to at least 925 nuclei per probe set. **(B)** For comparison, the same smFISH experiment was performed on chemically fixed cells. One representative cell is shown.

SmFISH dots detected for the different mRNAs ranged from, on average, 4.7 (*α-TUBULIN*) to 0.2 (*HEL67/DBP1*) per nucleus. The number of visible nuclei was used as a means of calibration to enable comparison between different slides and probe sets: as nuclei are subcellular structures of defined shape that occupy ~12% of the cell volume ^26^ they can serve as a proxy for cell numbers. Calibrating on *TUBULIN*, we calculated the number of mRNA molecules for the other four mRNAs from the smFISH data with the assumption that an increase/decrease in the number of probe pair recognition sites proportionally changes the number of RNA dots. Thus, an mRNA with 20 probe pair recognition sites is expected to produce double the number of dots in comparison to an equally abundant mRNA that is only recognised by 10 probe pairs (compare Figure 2B). Note that this is a simplification, as the true relationship between probe pair recognition sites and mRNA dots is complex and depends on many factors such as RNA structure (see below and discussion). One further correction was done for the *EP1* mRNA to compensate for the fact that this mRNA is part of a gene family and a subset of the probe pairs also recognises other family members (details in Figure legend of Figure 3A).

For *EP1* and *RPL7a*, the numbers calculated from the smFISH data were 582 and 268 mRNAs per cells, respectively, and thus almost identical to the mRNA numbers obtained from RNA sequencing (572 and 274, respectively). For *HEL67/DBP1*, we obtained a 1.6-fold higher number of mRNA molecules from FISH (33) in comparison to RNA sequencing (21). For *GB4*, smFISH data indicate 9.8 mRNAs per cell, in good agreement to smFISH data on chemically fixed cells (Supplementary Figure S4, Figure 3B and reference ^2^) while only 3 mRNAs per cell were measured by RNA sequencing. This discrepancy can be explained by a feature unique to very large mRNAs: we have previously shown that >50% of *GB4* is in the process of 5’-3’ decay (Supplementary Figure S4 and reference ^2^). These mRNA molecules in degradation are missed by typical RNA sequencing approaches as they lack both the miniexon and the poly(A) tail used for mRNA enrichment, but they are detected by smFISH. A further significant fraction of very large mRNAs is in the process of being transcribed (unknown for *GB4*, but 31% for the *FUTSCH* mRNA, which is of similar size to *GB4* ^2^), and these will be missed in any sequencing reaction based on poly(A) enrichment. Note that for average sized mRNAs, the fraction of decay intermediates is small and can be neglected ^2^.

The poly(A) probe (red dots in Figure 3) detected, on average, 16.6 dots per nucleus. The number of probe recognition sites for the poly(A) probe is difficult to predict. In contrast to all other Affymetrix probes, the poly(A) probe consists of a single oligo (of 29 Ts) modified to give the branched DNA amplification signal without the need of a second oligo binding next to it: two oligo(dT) probes binding exactly next to each other on a stretch of poly(A) is considered too unlikely. An average trypanosome poly(A) tail is about 100 nucleotides long ^27^, providing theoretical space for about 3 oligos. The real number of probe recognition sites is lower as oligos will bind randomly to the poly(A) tail, leaving gaps too small to be filled by probe pairs. Assuming 61,000 mRNA molecules per cell ^25^, the experimentally determined number of binding sites for the poly(A) probe per mRNA molecule is ~1.

These considerations allow calculating how much the number of detectable mRNA molecules is reduced by the 2D probing on the resin surface, in comparison to the 3D probing in chemically fixed cells (reduction factor). For the three averaged sized mRNAs probed with the standard amount of 20 probe pairs (*α-TUBULIN*, *RPL7a*, *HEL67/DBP1*) the reduction factor is 173.2±0.4. This factor changes reverse-proportionally to the number of probe pair recognition sites and is 7.4 for the large *GB4* mRNA and 3681 for the poly(A) probe.

In conclusion, LR White smFISH allows quantification of both low abundant and highly abundant mRNAs with accuracy comparable to RNA sequencing.

### Detection of mRNAs within dense subcellular structures (RNA granules)

One motivation for the development of LR White smFISH was the need to quantify mRNAs within protein-dense structures, in particular within RNA granules. There are two ways to calculate the fraction of an individual mRNA in granules by smFISH: either by measuring mRNA fluorescence inside and outside of the granules, or by counting individual mRNA molecules inside and outside of the granules. In chemically fixed trypanosomes we found both methods lacking accuracy: quantification via fluorescence requires that each mRNA molecule reveals approximately equal fluorescence, at least within the same cell. This is not the case. In particular mRNAs within RNA granules tend to be less fluorescent, for steric reasons (Figure 4A, triangle). Quantification via counting mRNA molecules requires that individual mRNAs can be distinguished, but this is often not possible because mRNAs are too abundant (Figure 4A, asterisk) or because more than one molecule accumulates in one granule (Figure 4A, arrow). For example, we found by biochemical fractionation mRNAs encoding ribosomal proteins underrepresented in stress granules, but we were unable to determine whether these mRNAs were just reduced or fully absent from granules ^6^ (Figure 4A, asterisk).

**Figure 4:**
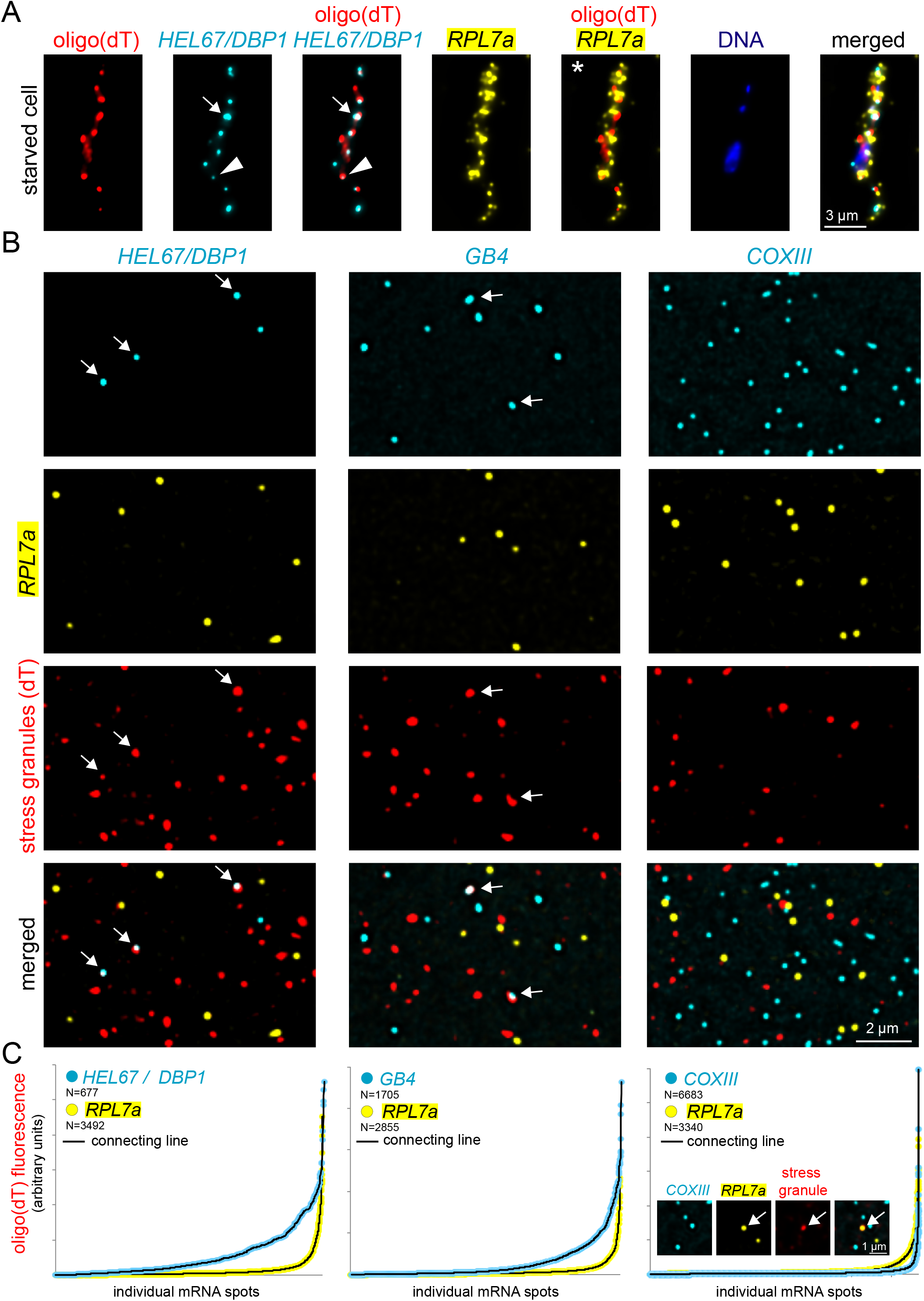
Detection of mRNAs in stress granules. **(A)** Starved trypanosomes were chemically fixed and probed for stress granules using an oligo(dT) probe (red) and probes for two specific mRNAs (*HEL67/DBP1* shown in cyan and *RPL7a* shown in yellow). For *HEL67/DBP1*, a huge variance in the intensity of the different mRNA dots is observed. Often, fluorescence within a granule is low (triangle). Strong, non-spherical signals indicate accumulation of more than one mRNA molecule in a granule (arrow). *RPL7a* was shown to be underrepresented from RNA granules ^6^, but its abundance prevents to distinguish whether it is fully excluded from granules or whether a small percentage is granule-localised (asterisk). **(B)** Samples of LR White embedded starved trypanosomes were probed for stress granules using an oligo(dT) probe (red). In addition, samples were incubated with Affymetrix probe sets for *RPL7a* (shown in yellow) and either *HEL67/DBP1*, *GB4* or *COXIII* (shown in cyan). Representative images are shown (projection of a Z-stack of 5 slices with 140 nm distance). **(C)** Individual mRNA dots were identified (find maxima function of FIJI) and the red fluorescence (oligo(dT) fluorescence = stress granule localisation) was measured in each dot. All mRNA dots are shown sorted according to their oligo(dT) fluorescence. The inlet (right graph) shows a rare example of a colocalisation between *RPL7a* and stress granules.

We anticipated that probing LR White embedded samples should resolve these problems. We starved trypanosomes by incubation in PBS for two hours to induce the formation of starvation stress granules ^6,8^ and embedded cells in LR White. For stress granule detection we used oligo(dT) coupled to a red fluorophore rather than branched DNA technology (the poly(A) probe set proved not suitable for this purpose). Stress granules had the expected variance in size and shape that we and others have previously observed (e.g. ^6^) (Figure 4B) and were absent on LR White embedded samples of non-stressed trypanosomes (not shown). In a second colour (shown in yellow) slides were hybridised with an *RPL7a* probe set; a representative of granule underrepresented mRNAs ^6^. In a third colour (shown in cyan), we probed for several control RNAs: *Hel67/DBP1*, *GB4* or unedited *COXIII*. *Hel67/DBP1* and *GB4* are mRNAs that localise to stress granules to a high percentage ^6^ (positive controls). *COXIII* is a mitochondrial mRNA (more details below) and should therefore not localise to cytoplasmic stress granules (negative control).

As expected, we observed frequent colocalisation of *HEL67/DBP1* and *GB4* mRNA molecules with stress granules (Figure 4B and C). Unlike in chemically fixed cells, granule-localised mRNA dots were of bright fluorescence (arrows Figure 4B), showing that the problem of probe accessibility has been solved. To quantify stress granule localisation, we measured oligo(dT) fluorescence in the maxima of the mRNAs dots and plotted the sorted oligo(dT) values (Figure 4C). For the mRNAs used in this experiment, stress granule localisation was highest for *HEL67/DBP1*, followed by *GB4*, *RPL7a* and *COXIII*. Granule localisation of *RPL7a* was significantly reduced in comparison to *HEL67/DBP1* and *GB4*, consistent with earlier findings ^6^; however, it was still higher than apparent granule localisation of *COXIII* and a very small fraction of *RPL7a* molecules was detectable in stress granules (Figure 4B and C). These data show that stress granule localisation of *RPL7a* is reduced, but not completely prevented.

### Monitoring mRNA compactness

The findings that only a small subset of probe pair recognition sites are accessible for probing each mRNA (Figure 2), while, at the same time, probe access has become independent of cellular environments (Figure 4B) prompted us to investigate the question, whether the method can be employed to monitor mRNA tertiary structure. We reasoned that mRNAs with compact structures should, on average, expose more probe pair recognition sites to the surface of the resin in comparison to mRNAs with extended structures, and hence, should have a higher fluorescence signal (Figure 5A and B). It is well established that mRNAs within stress granules are more compact than mRNAs outside of granules (e.g. ^28,29^). We therefore reused the stress granule data from Figure 4 to investigate whether LR White smFISH can identify differences in signal intensity between mRNAs in granules and mRNAs outside of granules. The mRNA dots of the two mRNAs *HEL67/DBP1* and *GB4* were divided into three groups of equal numbers, corresponding to low oligo(dT) fluorescence (no granule localisation), high oligo(dT) fluorescence (granule localisation) and intermediate oligo(dT) fluorescence. For each group, we measured the maximum fluorescence of each mRNA dot. We found granule-localised *HEL67/DBP1* and *GB4* mRNA dots 1.4- and 2.1-fold more fluorescent than mRNA dots outside of granules, respectively, and these differences were highly significant (t-TEST<0.005) (Figure 5C). The intermediate group also showed a significant increase in fluorescence in comparison to the no-granule group. We can exclude that this increase in fluorescence results from detecting more than one RNA molecule within the same granule because the vast majority of granules does not contain *HEL67/DBP1* or *GB4* dots (Figure 4B). The large range of fluorescence intensities within all groups of mRNAs is expected and reflects the random orientation of the molecules in 3D, which also causes variances in the number of surface-exposed probe pair recognition sites.

**Figure 5:**
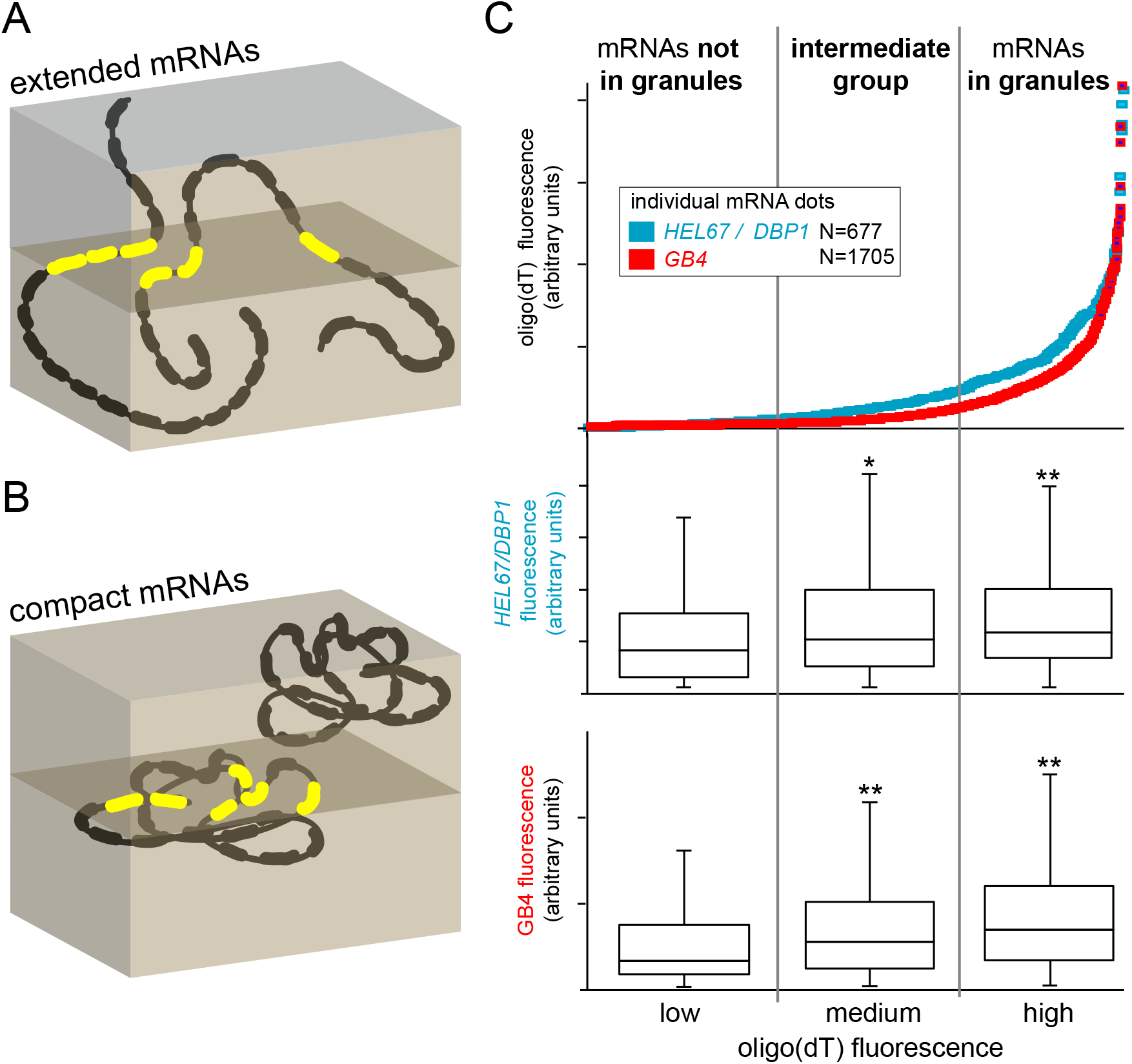
Monitoring mRNA structure. **(A)** 3D schematics of two extended mRNA molecules, each with 20 probe pair recognition sites. In this example, each mRNA has three probe pair recognition sites available for probing at the surface of the cut (red). **(B)** 3D schematics of two compact mRNA molecules, each with 20 probe pair recognition sites. In this example, one molecule has six probe pair recognition sites available for probing in the plane of cut (red), while the second molecule is not detected. **(C)** LR White embedded, starved trypanosomes were probed for *HEL67/DBP1* or *GB4* as well as for stress granules (oligo(dT)). mRNA dots were sorted according to the oligo(dT) fluorescence in their maxima (data are from Figure 4C) and divided into three groups: mRNA dots with low oligo(dT) fluorescence (not in granules), medium oligo(dT) fluorescence (granule localisation not certain) and high oligo(dT) fluorescence (localised in granules). For each group, the fluorescence maxima of the *HEL67/DBP1* or *GB4* dots were measured, and the data are shown as a box plot (waist is median; box is interquartile range (IQR); whiskers are ±1.5 IQR). Significant (t-TEST<0.05) and highly significant (t-TEST<0.005) differences to the dots with low stress granule localisation are indicated with * and **, respectively.

The data establish LR White smFISH as a novel tool to detect differences in mRNA tertiary structures between different populations of RNAs.

### Combination with immunofluorescence and electron microscopy

LR White smFISH can be easily combined with immunofluorescence detection and with electron microscopy. As a proof of principle, we probed LR White embedded BSF trypanosomes with Affymetrix probe sets for *VSG* mRNAs (shown in yellow), followed by incubation with an antibody for the VSG protein (red). After imaging, samples were contrasted, imaged by scanning electron microscopy, and the images were correlated. An example image is shown in Figure 6A. VSG, with 1*10^7^ copies per cell ^30,31^, is the most abundant protein in trypanosomes. A strong antibody signal was observed at the cell surface, where 90% of the VSG is localised, as well as in some intracellular places that are involved in VSG trafficking and recycling. For 3D imaging, 41 slices of LR White embedded trypanosomes, with 100 nm thickness, were probed for *VSG* mRNA, imaged by scanning electron microscopy and assembled (Figure 6B and Supplementary Movie S1). The same mRNA dot was never observed on two adjacent slices, confirming that the ‘depths’ of probe penetration into the LR White resin is <<100 nm. This is in agreement with the fact that the standard ‘reduction factor’ of the smFISH (compare Figure 3) is with 173 much larger than the average number of 20-30 100-nm slices each individual cell is covered by.

**Figure 6:**
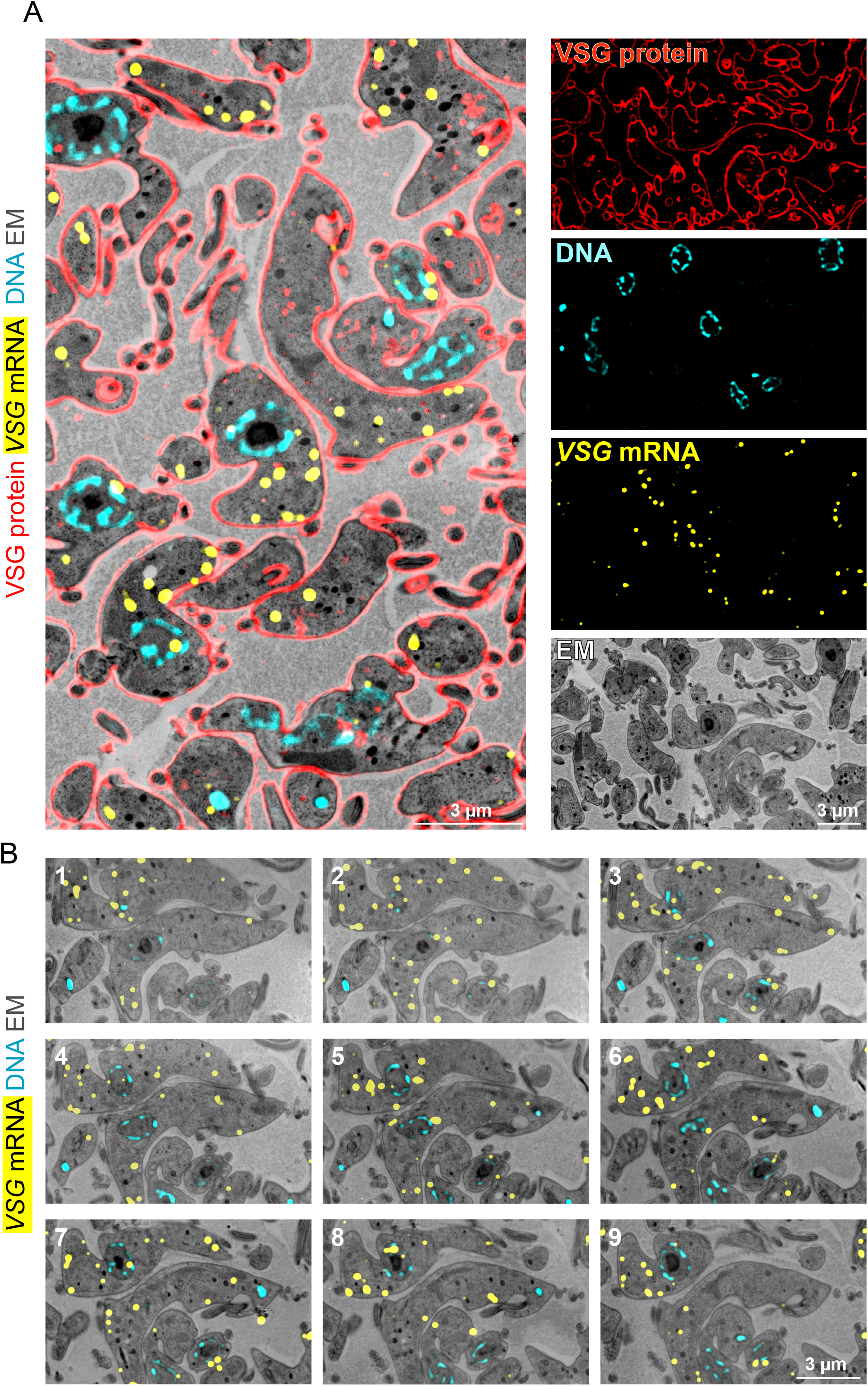
Combination of LR White smFISH with immunofluorescence and electron tomography. **(A)** LR White embedded BSF trypanosomes were probed for *VSG* mRNA (shown in yellow, imaged by standard fluorescence microscopy), followed by antibody incubation with anti-GFP (red, imaged by structured illumination microscopy) and DNA staining with DAPI (cyan). After imaging, samples were contrasted and imaged on a scanning electron microscope. Correlated images and individual images are shown. **(B)** Samples from (A) were cut into 41 sections, with 100 nm thickness. All slices were probed for *VSG* mRNA, stained with DAPI for DNA and imaged by fluorescence microscopy. Samples were contrasted, imaged by scanning electron microscopy and assembled to a stack. Nine images of the stack are shown; for the entire stack see Supplementary movie S1.

Having established LR White smFISH as a robust method to analyse various aspects of mRNAs on single molecule level, below we present two applications of the method related to trypanosome RNA biology.

### Application to trypanosome RNA biology I: RNA editing

Trypanosomes edit some of their mitochondrion-encoded mRNAs by additions and deletions of uridines ^17^. In extreme cases mRNAs are pan-edited, i.e. their sequences are dramatically altered by frequent insertion (and occasional deletion) of uridine residues along their entire length, which can result in a doubling in mRNA size. RNA editing is achieved by a complex enzymatic machinery together with guide RNAs, the latter being mainly encoded by so-called minicircle DNA molecules of the kinetoplast. RNA editing proceeds from the 3’ to the 5’ end of an mRNA ^32^. RNA sequencing of steady-state populations of two small pan-edited mRNAs (*RPS12* and *ND7*) revealed that the majority of all molecules are RNA editing intermediates, 14% and 31% are unedited RNAs, respectively, and only a very small fraction (<6%) are fully edited RNAs ^33^. The cellular localisation of RNA editing is not clear, but several editing enzymes have been shown to be localised to distinct antipodal nodes of the kinetoplast or to the kinetoplast in general, although broad mitochondrial localisation was observed as well ^34–36^. Moreover, at least two ribosomal proteins locate to kinetoplast associated nodes close to the RNA editing enzymes, suggesting that transcription, editing and translation could be spatially linked ^34,35^. To the best of our knowledge, accurate localisations of mitochondrion-encoded mRNAs have not been reported.

To investigate localisation and abundance of edited and unedited mitochondrial mRNAs, we chose mRNAs encoding ATP synthase F_O_ subunit 6 (A6) and cytochrome *c* oxidase subunit III (COXIII). The F_O_F_1_ ATP synthase is expressed in both PCF and BSF *T. brucei* ^37^. While cytochrome *c* oxidase is only expressed in the PCF stage ^38^, complete editing of the *COXIII* mRNA in the BSF stage has been reported in some studies but not in others and may depend on the cell line under study ^39,40^. Both mRNAs are pan-edited ^39,41^ and of sufficiently large size to allow the design of specific probe sets against both the edited and unedited version, despite the presence of low complexity sequences that in particular edited mRNAs in particular are enriched for (Supplementary Figure S5, Figure 7A).

**Figure 7:**
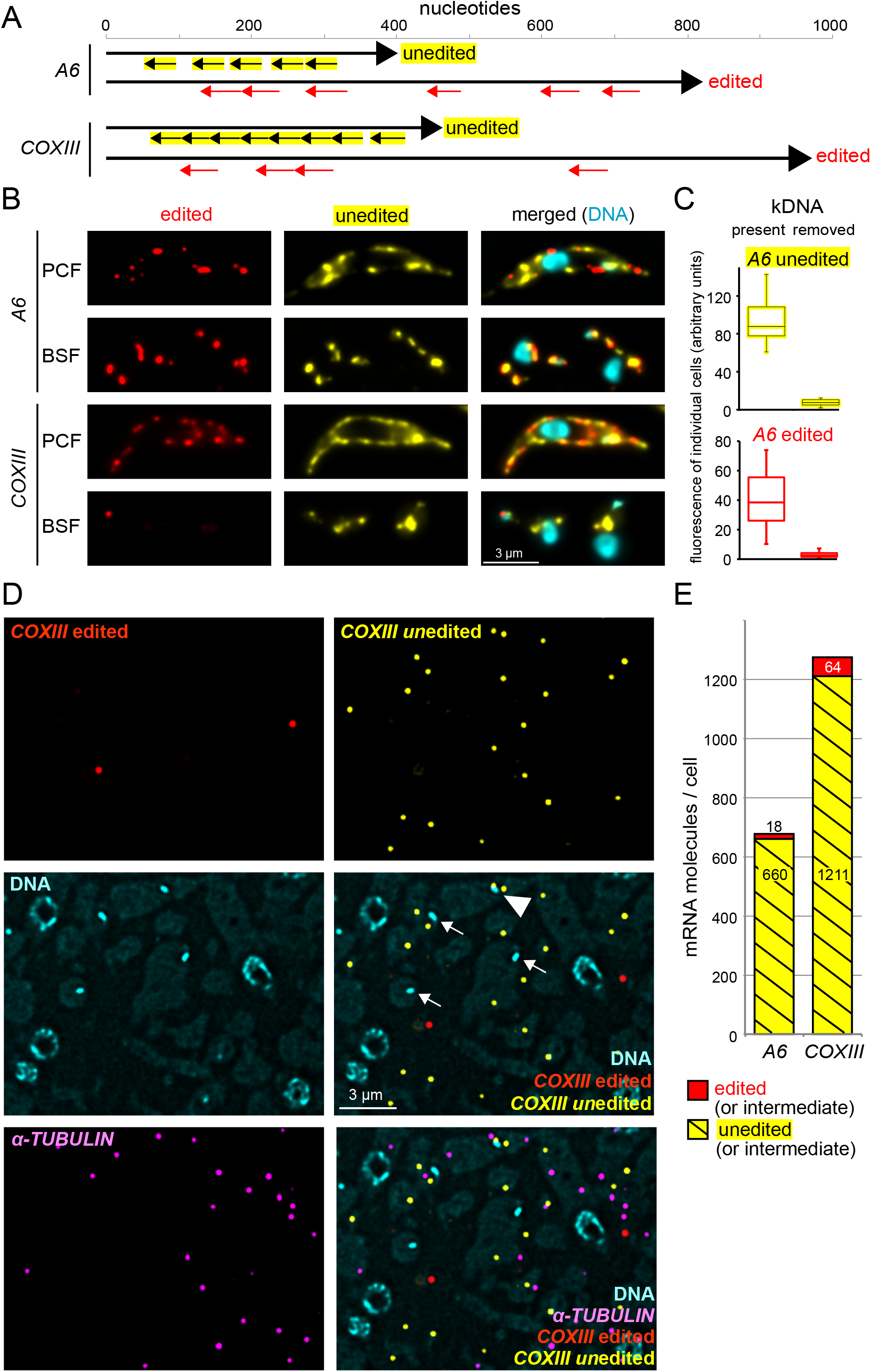
Detection and localisation of edited and unedited mitochondrial RNAs. **(A)** Schematics of the *A6* and *COXIII* edited and unedited mRNAs showing the positions of the probe pair binding sites. Edited mRNAs have regions of low sequence complexity due to the large number of uridine insertions, and large sections of the mRNA had to be omitted from probing or even blocked with special blocking oligos (further details on the probes are in Supplementary Table S1 and Supplementary Figure S5). **(B)** Chemically fixed BSF and PCF cells were probed as indicated. One representative cell is shown (projection of a deconvolved Z-stack with 50 slices a 140 nm). For further images, see Supplementary Figure S6, S7A and S8A. **(C)** To obtain additional evidence for the specificity of the probe sets, we used chemically fixed BSF cells that lacked kDNA and therefore cannot produce mitochondrial transcripts. Such akinetoplastic trypanosomes can be produced by treatment with 10 nM ethidium bromide for three days ^42,43^ and, if cells carry the compensating mutation L262P in their ATPase gamma subunit, they can survive this treatment ^44,45^. Control cells and akinetoplastic trypanosomes were probed for edited and unedited *A6*, and as a control, for the cytoplasmic mRNA *FUTSCH*. We observed a mild reduction in the number of *FUTSCH* dots per cell (from 1.7±1 to 0.9±0.9; N>220), possibly caused by the minor effects 10 nM ethidium bromide have on nuclear DNA. To compensate for this effect, we only included cells with exactly two FUTSCH dots each in the further analysis. The fluorescence of edited and unedited *A6* signal was measured from ten individual cells with exactly two FUTSCH dots and the (background-corrected) data are presented as boxplot (waist is median; box is interquartile range (IQR); whiskers are ±1.5 IQR). Akinetoplastic cells have, on average, 13 and 14 times less fluorescence of unedited and edited *A6*, respectively, than control cells. Microscopy images are shown in Supplementary Figure S7. The corresponding dataset for *COXIII* is shown in Supplementary Figure S8. **(D)** LR White embedded PCF cells were probed as indicated. One representative image is shown as deconvolved Z-stack projection (5 slices a 140 nm). Most mRNA molecules are not close to the kinetoplasts (which are marked by arrows). One rare exception of an unedited *COXIII* molecule close to a kinetoplast is marked with a triangle. Images of samples probed for *A6* are shown in Supplementary Figure S9. **(E)** The numbers of mRNA molecules per cell for both edited and unedited *A6* and *COXIII* mRNAs were calculated as described above (Figure 3). The total number of nuclei used for quantification was 5841 and 2909 for *A6* and *COXIII*, respectively.

First, we used the probes on chemically fixed cells of both the BSF and the PCF life cycle stage. For the PCF trypanosomes, which have a branched mitochondrion that is active in oxidative phosphorylation, a strong smFISH staining signal was observed for both unedited and edited *A6* and *COXIII* (Figure 7B shows representative single cells; Supplementary Figure S6 shows populations of cells). The signal distribution clearly differed from a cytoplasmic localisation signal by being restricted to thin structures close to the cellular periphery, consistent with the structure of the PCF mitochondrion. The mRNAs were too abundant to allow detection of single mRNA molecules. For BSF trypanosomes, which have a structurally more simple, tubular mitochondrion inactive in oxidative phosphorylation, the smFISH signal was still very abundant, with the exception of the edited *COXIII* mRNA that was detectable as single molecules in a few cells and undetectable in most cells (Figure 7B and Supplementary Figure S7, S8A). To obtain additional evidence for the specificity of the probe sets, BSF cells were experimentally prevented from producing mitochondrial transcripts by removing the kDNA (the DNA of the kinetoplast) through ethidium bromide treatment ^42–45^. Both the edited and the unedited *A6* signal were at least 13 times reduced, while a cytoplasmic control mRNA was only mildly affected (less than 2-fold reduction in dot number) (Figure 7C and Supplementary Figure S7). Similar results were obtained for unedited *COXIII* (Supplementary Figure S8).

Next, the experiment was performed on LR White embedded samples of PCF trypanosomes, using *α-TUBULIN* as marker for cytoplasmic RNA (Figure 7D and Supplementary Figure S9). Single mRNA molecules were resolved and allowed the quantification of unedited mRNAs (yellow dots) and edited mRNAs (red dots) mRNA sequences. With, on average, 678 molecules per cell, *A6* was as abundant as *α-TUBULIN*. *COXIII* was with 1275 molecules per cell twice as abundant as *α-TUBULIN*. For both mRNAs, the edited fraction contributed only 5% or less (Figure 7E), consistent with published data on *A6* ^32^. There were only very few double-coloured dots (one rare example is visible in Supplementary Figure S9), which would be indicative of mRNAs in the editing process. However, LR White smFISH is not suitable for such direct detection of RNA editing intermediates by double staining, because the expected number of probe sites on the resin’s surface is too small (compare Figure 2A). Notably, red and yellow dots do not indicate exclusively fully edited vs. fully unedited mRNAs, respectively: in both cases, it is possible that the mRNA is an editing intermediate with the ‘other part’ not accessible to the probe. The massive excess in unedited RNA sequences, however, indicates a large proportion of unedited RNA or RNA editing intermediates at the beginning of editing (see discussion). Importantly, the vast majority of both unedited and edited mRNA did not localise close to the kinetoplast (arrow in Figure 7D, Supplementary Figure S9), with only rare exceptions (triangle in Figure 7D).

The LR White smFISH has allowed detecting, quantifying and localising both edited and unedited mitochondrial mRNAs for the first time. Neither edited nor unedited mRNAs showed any obvious enrichment to a certain sub-region of the mitochondrion, including the kinetoplast region, providing no evidence for a spatial coupling of transcription and uridine insertion/deletion editing.

### Application to trypanosome RNA biology II: the SL RNA

The SL RNA is a trypanosome-specific 139 nucleotide long RNA that is used in the trans-splicing reaction and consists of the capped miniexon (nucleotides 1-39) and the SL RNA intron (nucleotides 40-139) (Figure 8A). The capped miniexon is transferred to every mRNA. It is not clear, whether the SL RNA localises mainly to the nucleus or also to the cytoplasm, as data from standard FISH experiments are inconsistent ^22,46,47^. Even though the trans-splicing reaction takes place in the nucleus, there is evidence that some SL RNA maturation steps may take place in the cytoplasm ^46,48,49^. Accurate localisation of the SL RNA is therefore essential.

**Figure 8:**
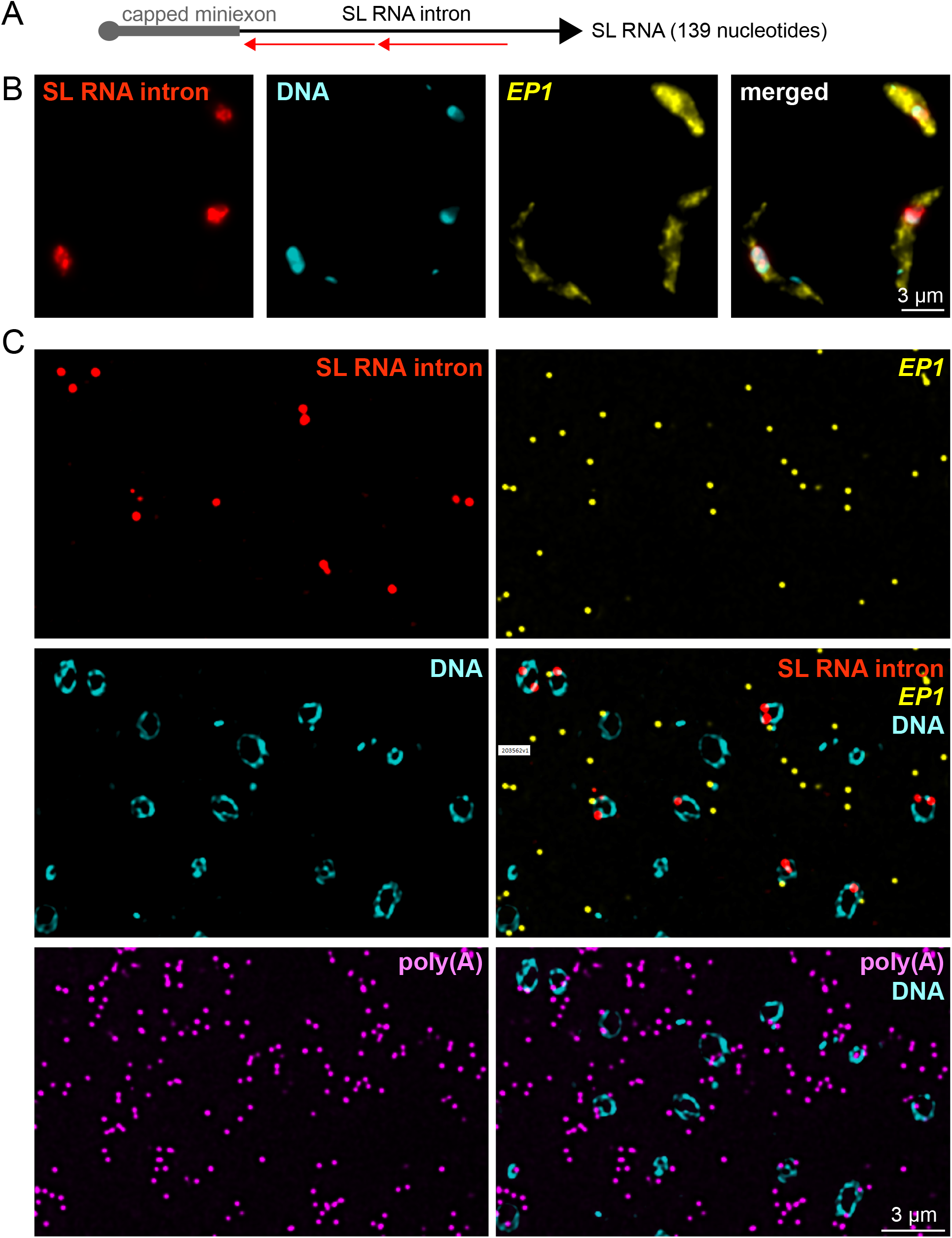
Localisation and quantification of SL RNA. **(A)** Schematics of the SL RNA with binding sites for the two probe pairs. **(B and C)** Chemically fixed (B) and LR White embedded (C) PCF trypanosomes were probed as indicated. One representative image is shown as projection of a deconvolved Z-stack with 50 and 5 images per 140 nm slice for panels B and C, respectively.

To visualise the SL RNA, we designed two probe pairs antisense to the SL RNA intron (Figure 8A). Probing for the intron can be considered specific to the detection of the unspliced SL RNA, as the amount of SL RNA splicing product is very small in comparison to the full length SL RNA and can be neglected ^50^. Again, we probed both chemically fixed cells and slices of LR White embedded samples. Both methods detected the majority of SL RNA in the nucleus, while control mRNAs (EP1, polyadenylated RNA) were mainly in the cytoplasm (Figure 8B and C, and Supplementary Figure S10). LR White smFISH provided quantitative data: more than 90% of all SL RNA dots were unequivocally within the nucleus. The calculated average number of SL RNA molecules per cell was 1100 and thus slightly higher than *α-TUBULIN* (810), but 55-fold less abundant than total mRNAs (61,000) ^25^. Within the nucleus, the number of polyadenylated mRNA molecules was in 3.8-fold excess to the number of SL RNA molecules (Supplementary Figure S10).

## DISCUSSION

With LR White smFISH we present a method that allows the simultaneous measurement of localisation, abundance and compactness for up to four RNAs in parallel, independent of subcellular localisation. Moreover, the method is directly applicable to any biological material (any type of cells or tissues), as probing is solely on the resin surface, avoiding any probe accessibility differences between or within samples. The method can be combined with electron microscopy and immunofluorescence, providing correlative information on subcellular RNA localisation.

We introduce LR White smFISH as a novel tool to probe for RNA compactness. Compact RNAs are densely packed within a small volume, while less compact RNAs are extended strings that are elongated at least in one dimension. While structural RNAs (tRNAs, rRNAs) have a defined compactness, mRNAs can exist in both compact and extended states with switches between the states being mediated by RNA binding proteins, changes in base-pairing, temperature or any combination of these. In LR White smFISH, the fluorescence of an RNA dot directly reflects the number of surface-exposed probe pairs that is the larger the more compact an mRNA molecule is (Figure 5A-B). As a proof of principle, we show on two examples that stress granule localised mRNAs are, on average, more compact than mRNAs outside of granules. mRNA compactness has been previously determined by smFISH via probing the 5’ and 3’ ends of very large RNA molecules in two different colours: the distance between these two dots reflects mRNA compactness ^28,29^. LR White smFISH has two advantages in comparison with the double-colour probing: (1) it works with average-sized RNAs on a standard light microscope (as long as multiple probe pairs are used) and (2) it uses only one colour per RNA, doubling the number of RNA species that can be detected simultaneously.

Are there limits for RNA detection and quantification? We have successfully quantified mRNAs ranging from 10 to 60,000 mRNA molecules/cell (note that the volume of a trypanosome cell is similar to the volume of a yeast cell; different detection ranges will apply to smaller or larger cell types). To reach this range we have adapted the number of probe pairs that recognise the target: total mRNAs (60,000/cell) were detected with only one probe per molecule, while the mRNA with lowest abundance (9/cell) had 468 probe pair recognition sites due to its repetitive nature. To detect an mRNA with 10 molecules/cell with the standard set of 20 probe pairs, one dot per 17 nuclei is expected, with our settings, this equals on average nine dots per microscopy image - still a feasible approach. The majority of RNA species have an abundance lying within the range studied here, for example snRNAs (we show the example of the SL RNA which serves as the snRNA for the U1 snRNP in trypanosomes) and also tRNAs (which we already have detected as single molecules by LR White smFISH, our own unpublished data). Only the highly abundant ribosomal RNAs may not resolve as single dots. Super-resolution microscopy could help to some extent. An alternative approach is to reduce the number of available probe pair recognition sites with a competing oligo (as done in Figure S2). Another approach is to use the total fluorescence signal rather than the number of dots for both localisation and quantification - the former limits of probe accessibility have been solved.

One limit in RNA quantification by LR White smFISH is that differences in RNA tertiary structures will affect quantification. More compact RNAs are less likely in the plane of cut than extended RNAs (compare Figure 5A and B) and thus appear less abundant in comparison. This limit can be overcome by probing with only one probe pair or by including the intensity of RNA dots into the calculation of abundance, using appropriate controls.

We have employed LR White smFISH to answer some outstanding questions of trypanosome RNA biology:

1. We show that *RPL7a* mRNA is not fully excluded from trypanosome starvation stress granules, but only largely underrepresented. The underrepresentation of mRNAs encoding ribosomal proteins from RNA granules is not specific to trypanosomes ^6^, but was also found when analysing purified P-bodies of mammalian cells ^51^; the reason for this phenomenon is unknown. Neither RNA sequencing of purified granules nor smFISH methods on chemically fixed cells can faithfully distinguish between absence and underrepresentation of an mRNA from granules and we propose to use LR White smFISH as an alternative to study functions and mechanisms of mRNA granule localisation.
2. We show that the SL RNA localises mostly to the nucleus. Localisation of the SL RNA was in the past examined by standard FISH, with two publications reporting a strong cytoplasmic staining in addition to the nuclear staining ^22,46^ and one reporting exclusive nuclear localisation ^47^. The likely reasons for these discrepancies are differences in background staining and probe accessibilities between methods. In branched DNA technology these problems are avoided. We could show predominantly nuclear localisation of the SL RNA, in both chemically fixed cells and on LR White embedded cells, consistent with ^47^. LR White smFISH did not only allow quantification of the SL RNA (1100 molecules / cell) but also showed the number of SL RNA molecules to be four times less than the number of nuclear-localised polyadenylated mRNAs. Given that each SL RNA produces one mRNA, this indicates that the half-life of the SL RNA is about four times shorter than the time, an average mRNA spends inside the nucleus after maturation. The half-life of the SL RNA is about four minutes ^52^, indicating that each mature mRNA spends on average 16 min inside the nucleus. While the notion that trans-splicing takes place in the nucleus is beyond any doubt, there is some evidence that certain SL RNA maturation steps take place in the cytoplasm. For example, a large fraction of the SL RNA can be found in cytoplasmic fractions after biochemical fractionation ^48,49^, and SL RNA export to the cytoplasm was suggested to depend on the nuclear exporter exportin (XPO1) ^46^. Our data do not fully exclude such a cytoplasmic maturation step, as LR White smFISH occasionally (<10%) detects cytoplasmic SL RNA. However, our recent data indicate that mRNA export in trypanosomes is not tightly regulated, if indeed it is regulated at all ^3^. An alternative explanation for the small amount of cytoplasmic SL RNA could therefore be the lack of an active mechanism that prevents the SL RNA from leaving the nucleus. Its predominant nuclear location could result from its short half-life. Furthermore, in biochemical fractionations small proteins preferentially leave the nucleus ^53^ and the same may be true for small RNAs, explaining the presence of the SL RNA in cytoplasmic fractions ^48,49^.
3. We have quantified and localised both edited and unedited mitochondrial mRNAs for the first time. With 678 and 1275 mRNA molecules per cell, respectively, *A6* and *COXIII* mRNAs were at least as abundant as *α-TUBULIN*, but only a very small fraction of these transcripts is fully or at least partially edited. This is consistent with earlier studies on *A6* ^32^ and with studies on two other pan-edited transcripts, *ND7* and *RPS12* ^32,33^. Higher fractions (up to 30%) of fully edited RNAs are only found in minimally edited mRNAs ^54^. With our experimental set-up, it is impossible to distinguish RNA editing intermediates from fully edited or fully unedited RNA species. Double-staining was too rare an event to allow meaningful conclusions. However, the massive excess in unedited RNA sequences in comparison to edited RNA sequences indicates that a large fraction is either unedited or an RNA intermediate at the beginning of editing. RNA editing proceeds from 3’ to 5’ end, but is not progressive and stops at several intrinsic pause sites (IPS) ^33,55,56^. One possible reason for the large percentage of *A6* mRNAs recognised by the unedited probe set could therefore be a strong IPS that was identified near the 3’end of the molecule ^55^, and a similar explanation might apply to *COXIII*. Although positions and numbers of IPSs have not been determined in a quantitative way for the large pan-edited mRNAs *COXIII* and *A6* that we investigated in this work, the available sequencing data on other edited mRNAs indicate that the percentage of edited RNAs decreases with the number of editing events ^33,54^. *COXIII* and *A6* have more editing events than the mRNAs sequenced so far providing an explanation for the large fraction of unedited sequences.

We found no significant enrichment of either edited or unedited RNA at a specific location within the mitochondrion, in particularly not at the kinetoplast. In the related parasite *Leishmania*, it was previously observed by immunofluorescence that components of RNA editing complexes as well as mitoribosomes localise to one or two antipodal nodes adjacent to the disk of the kDNA, indicating that transcription, editing and translation may be linked ^34,35,57^. A link between transcription and editing would require the unedited RNA to localise close to the kinetoplast, which is the site of transcription; and if translation really takes place at the inner-mitochondrial membrane close to the kinetoplast, the translated fraction of the edited RNA is expected to localise there too. Possible explanations for this apparent discrepancy are that the situation in *T. br*ucei could be different to *Leishmania* or that the two mRNAs investigated here are exceptions rather than typical. Importantly, however, a fraction of editing complexes in *Leishmania* was found to be dispersed throughout the mitochondrion, active vs. inactive complexes could not be distinguished in that study ^34,35^, and components of editing complexes localised as part of the genome-wide protein localisation project TrypTag ^36^ were variably found associated with the kinetoplast as well as dispersed throughout the mitochondrion.

In conclusion, we have established LR White smFISH as a method that provides information on mRNA localisation, abundance and structure in one single experiment, with high accuracy and importantly, independent of the biological material/organism. Furthermore, combination with other imaging methods, most notably electron microscopy tools, can fine-tunes localisation studies. The Affymetrix kit provides a maximum of 4 colours to be used simultaneously, but with multiplexing systems like MERFISH ^58^, which can also be combined with branched DNA technology ^59^, whole genomes can be probed for in the future.

## ACKNOWLEDGEMENTS

We are grateful for the help of Sebastian Markert and Sebastian Britz (Zentrale Abteilung für Mikroskopie, Biozentrum, Universität Würzburg, Würzburg, Germany) with structured illumination microscopy and correlative microscopy.

## SUPPLEMENTARY MATERIAL

**Supplementary Figure S1:** Specificity of the *RPL7a* probe set

**Supplementary Figure S2:** Specificity of the poly(A) signal

**Supplementary Figure S3:** Life cycle specific mRNAs are only detectable in the respective life cycle stage.

**Supplementary Figure S4:** Two-colour smFISH of *GB4* mRNA in chemically fixed cells

**Supplementary Figure S5:** Probe sets for edited and unedited *A6* and *COXIII* mRNAs.

**Supplementary Figure S6:** Chemically fixed PCF trypanosomes were probed for unedited or edited *A6* or *COXIII* mRNAs.

**Supplementary Figure S7:** Specificity test for the *A6* probe set using akinetoplastic trypanosomes

**Supplementary Figure S8:** Specificity test for the *COXIII* probe set using akinetoplastic trypanosomes

**Supplementary Figure S9:** LR White probing for *A6*

**Supplementary Figure S10:** Quantification of nuclear-localised SL RNA and poly(A)

**Supplementary Movie S1:**

41 slices of LR White embedded trypanosomes were probed for *VSG* mRNA, imaged by scanning electron microscopy and assembled. This video shows all 41 slices.

**Supplementary Table S1:**

This table contains detailed information about all Affymetrix probe sets.

**Supplementary Table S2:**

This table contains information on the calculations of how many dots of each colour/colour combination are expected in the two-colour competition assay (Figure 2C), in dependency of the number of probe pairs.

