## Supplementary Figures for "Parallel monitoring of mRNA abundance, localisation and compactness with correlative single molecule FISH on LR White embedded samples"

Figure S1

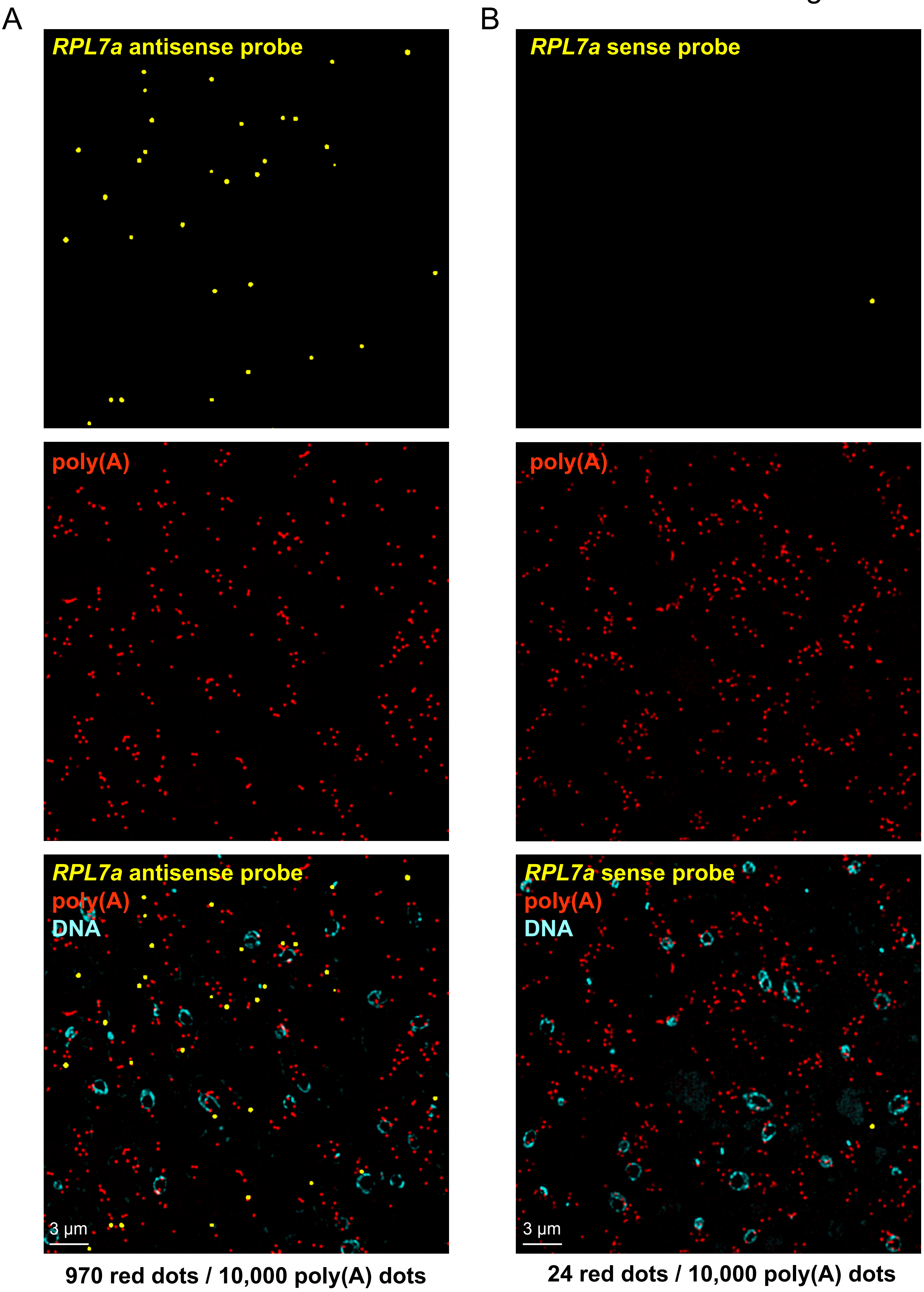


**Supplementary Figure S1: Specificity of the *RPL7a* probe set**
LR White embedded PCF trypanosomes were hybridised with a probe set antisense **(A)** or sense **(B)** to the *RPL7a* mRNA (yellow) and with a poly(A) probe set (red). DNA was detected with DAPI. The number of yellow dots per poly(A) dots was quantified for >20,000 poly(A) dots and is indicated below the images.

Figure S2

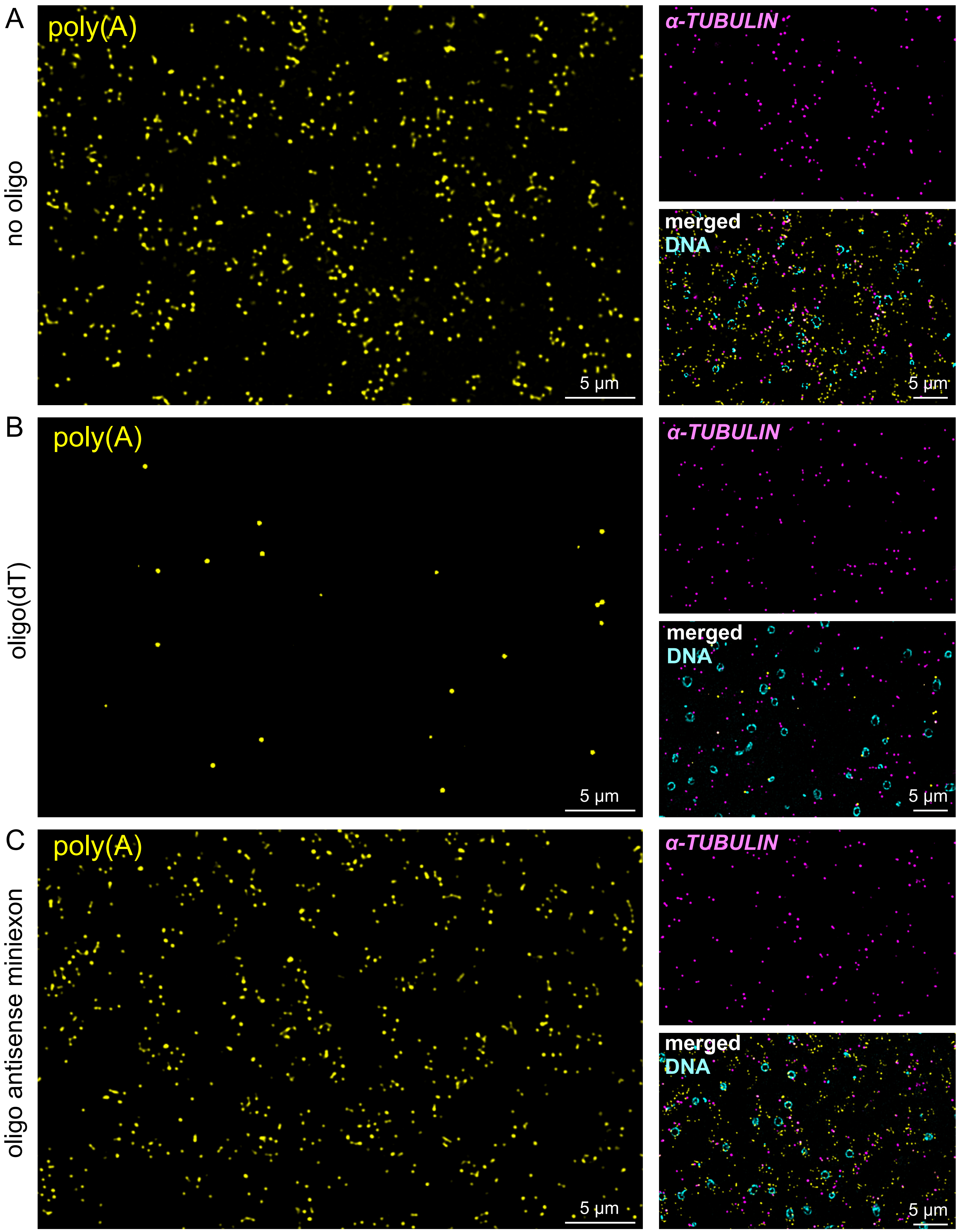


**Supplementary Figure S2: Specificity of the poly(A) signal**

smFISH was performed on LR White embedded PCF trypanosomes using probepairs antisense to poly(A) (yellow) and to α-*TUBULIN* (pink). The reaction was performed either in the absence of oligos **(A)**, in the presence of 2 µM oligo(dT) (30 nucleotides) **(B),** or in the presence of 2 µM of an oligo antisense to the miniexon sequence, 5´- CAATATAGTACAGAAACT-GTTCTAATAATAGCGTT -3´ (**C**). Deconvolved Z-stack images (sum slices of 5 stacks of 140 nm each) are shown. DNA was detected with DAPI.

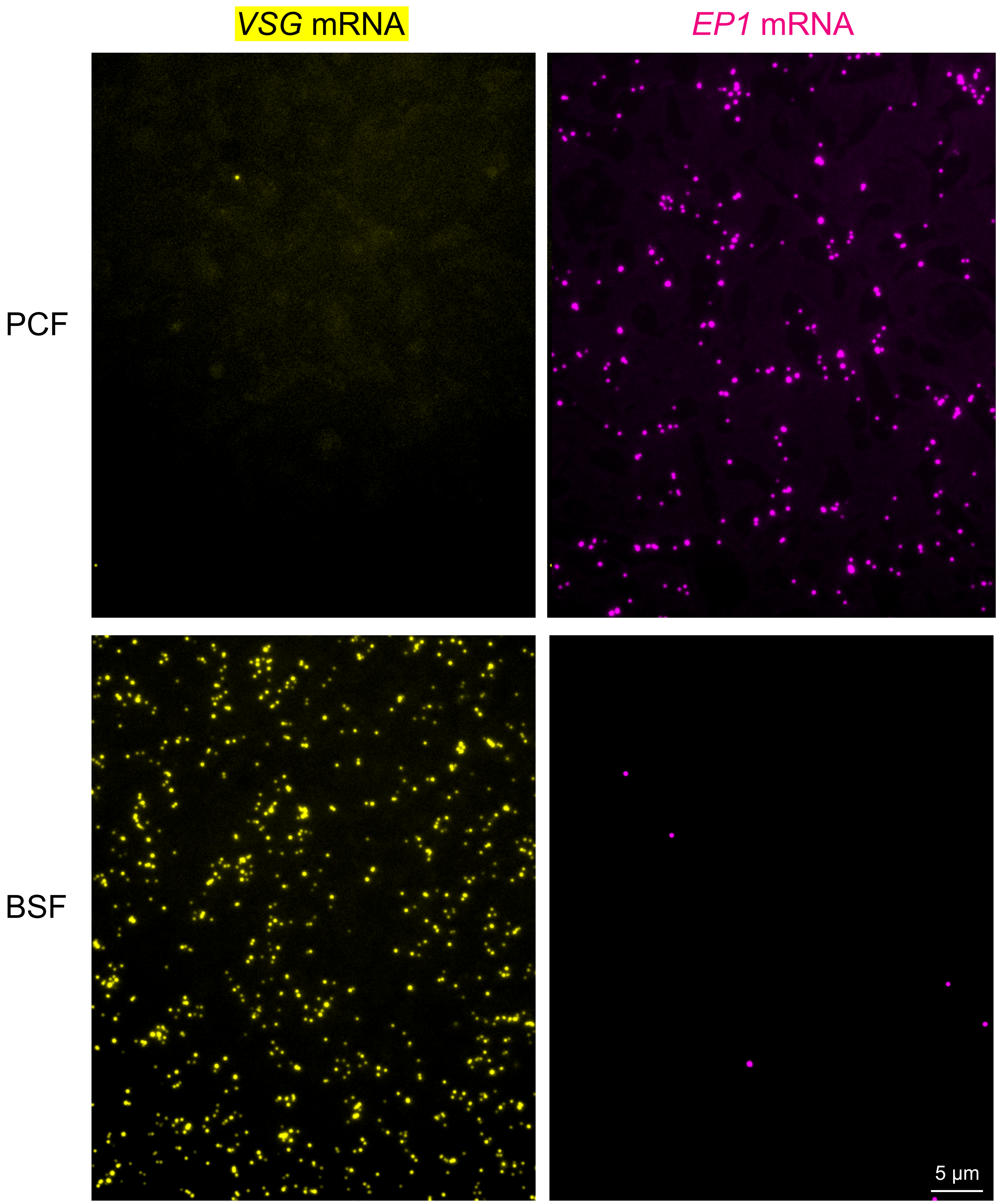


**Supplementary Figure S3: Life cycle specific mRNAs are only detectable in the respective life cycle stage.**

LR White embedded BSF and PCF trypanosomes were probed for the PCF-specific mRNA *EP1* (pink) and for the BSF specific mRNA *VSG* (yellow). In the BSF stage, 0.84% of mRNA dots were *EP1* (N=14,331). In the PCF stage, 0.36% of all mRNA dots were *VSG* (N=9,282). It remains unknown whether these few dots (*EP1* in BSF and *VSG* in PCF) reflect the true expression rates or whether these are due to non-specific staining.

### Figure S4

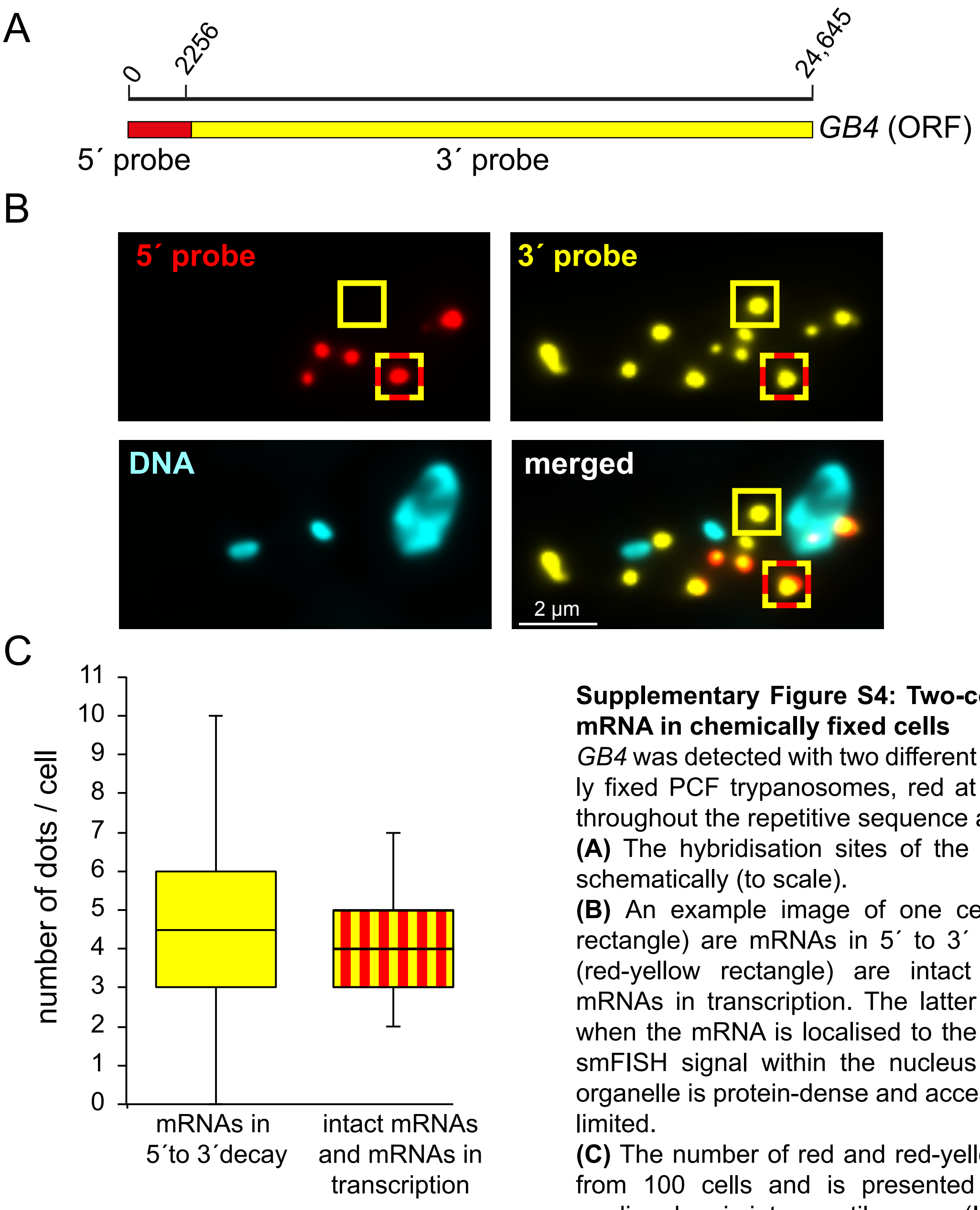


**Supplementary Figure S4: Two-colour smFISH of *GB4* mRNA in chemically fixed cells**

*GB4* was detected with two different probe sets in chemically fixed PCF trypanosomes, red at the 5´ end and yellow throughout the repetitive sequence at the 3´end.

**(A)** The hybridisation sites of the probe sets are shown schematically (to scale).

**(B)** An example image of one cell: yellow dots (yellow rectangle) are mRNAs in 5´ to 3´ decay, red-yellow dots (red-yellow rectangle) are intact mRNA molecules or mRNAs in transcription. The latter is in particularly likely when the mRNA is localised to the nucleus. Note that the smFISH signal within the nucleus is poor, because this organelle is protein-dense and accessibility of the probes is limited.

**(C)** The number of red and red-yellow dots was quantified from 100 cells and is presented as box-plot (waist is median; box is interquartile range (IQR); whiskers are ±1.5 IQR).

Figure S5

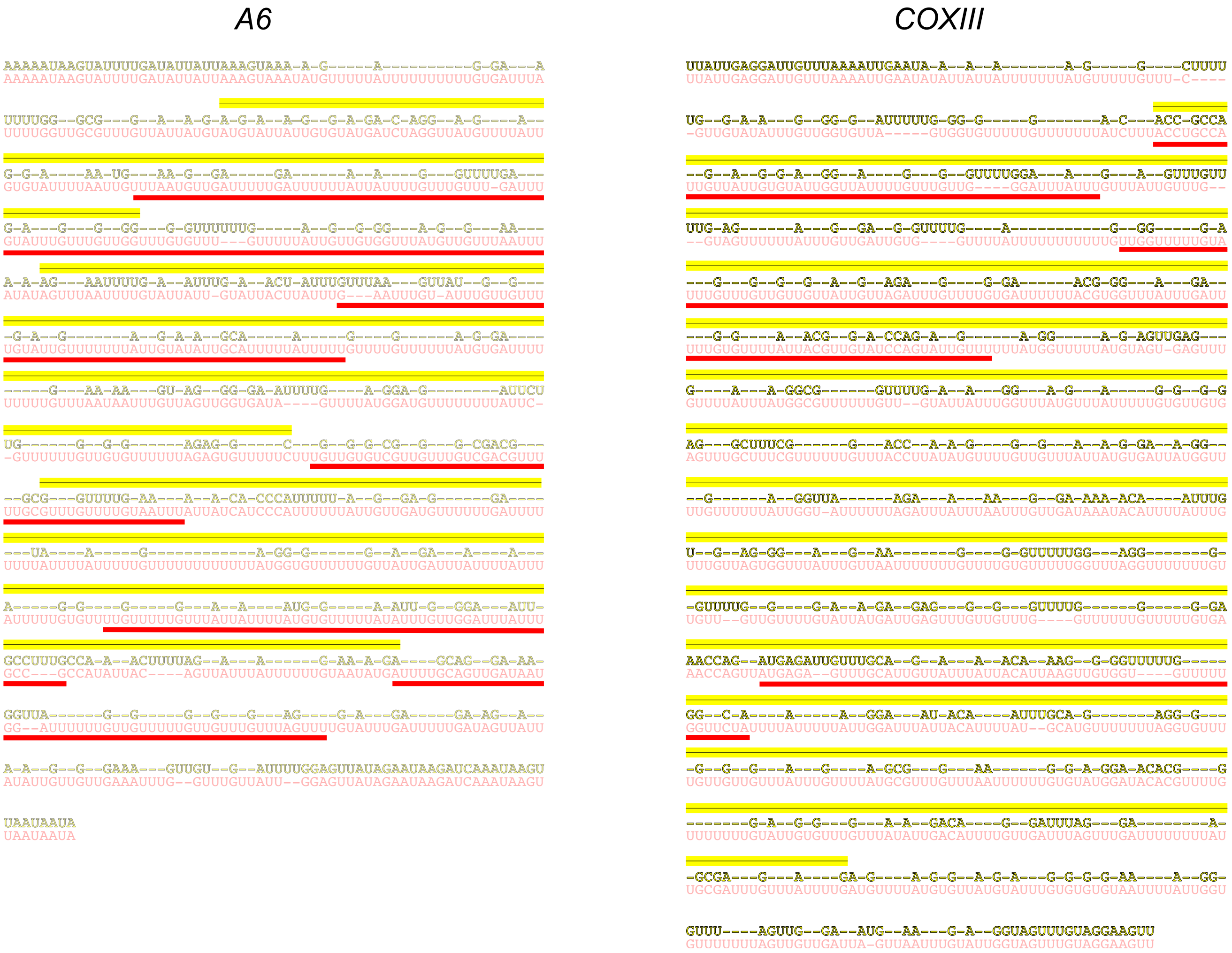


**Supplementary Figure S5: Probe sets for edited and unedited A6 and COXIII mRNAs.**
Edited (light red) and unedited (yellow) sequences are aligned for *A6* and *COXIII* mRNAs.The hybridisation sites for the smFISH probe pairs are indicated with red and yellow bars, for edited and unedited mRNA, respectively.

Figure S6

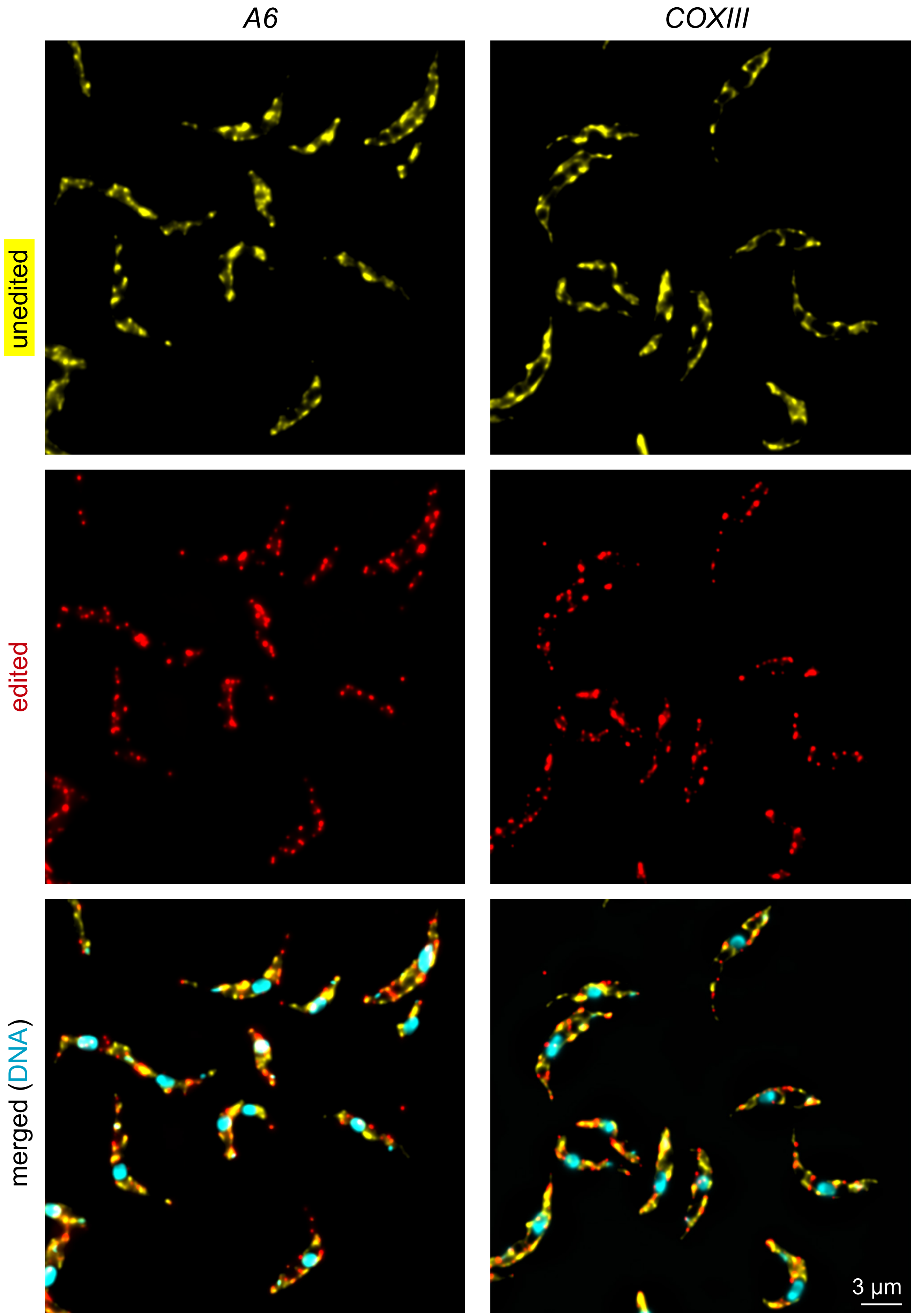


**Supplementary Figure S6:** Chemically fixed PCF trypanosomes were probed for unedited or edited *A6* or *COXIII* mRNAs. DNA was stained with DAPI. Projections (sum slices) of deconvolved Z-stack images (50 stacks a 140 nm) are shown.

### Figure S7

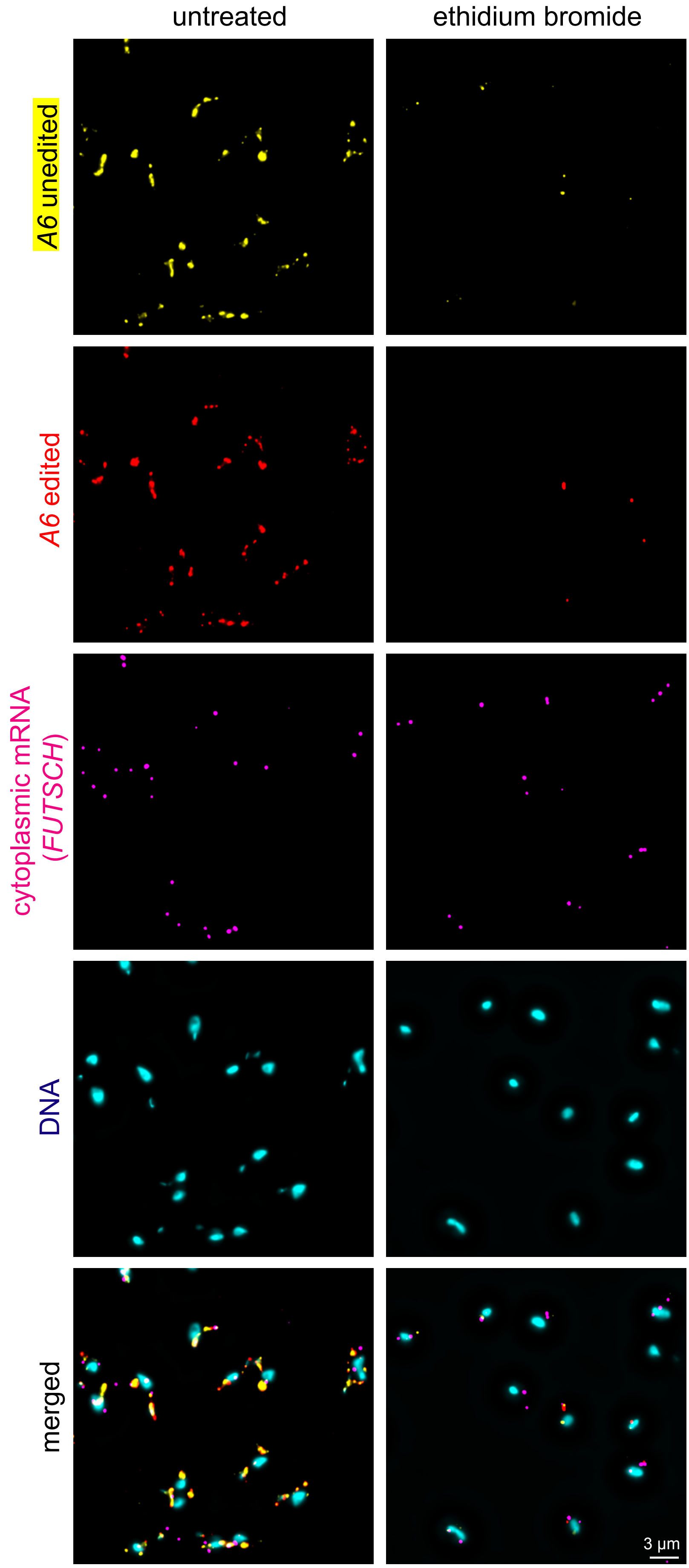


**Supplementary Figure S7: Specificity test for the *A6* probe set using akinetoplastic trypanosomes** BSF trypanosomes carrying a mutant of the ATPase gamma subunit (L262P) were treated with 10 nM ethidium bromide for three days to remove kDNA; untreated parasites served as control. Chemically fixed trypanosomes were probed via smFISH for edited and unedited *A6* mRNA as well as for *FUTSCH* (a cytoplasmic control mRNA).

Projections (sum slices) of deconvolved Z-stacks (50 slices a 140 nm) are shown. Note the lack of kinetoplasts in the ethidium bromide samples (DNA stain) as well as the massive reduction in fluorescence for both the edited and unedited *A6* mRNA. In contrast, for the *FUTSCH* control mRNA, we only observed a mild reduction in the number of dots per cell (from 1.7±1 to 0.9±0.9; N>220), possibly caused by the minor effects 10 nM ethidium bromide have on nuclear DNA. Quantitative data are shown in Figure 7C.

### Figure S8

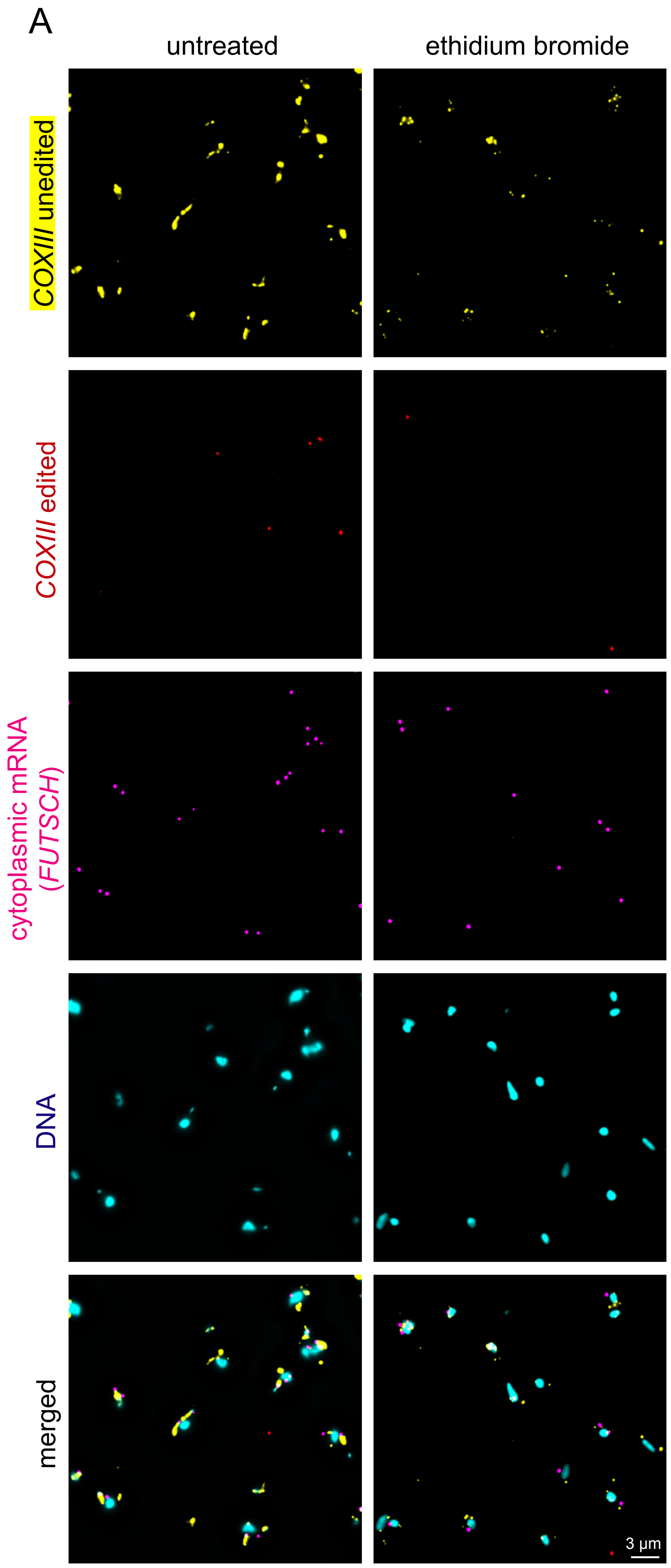


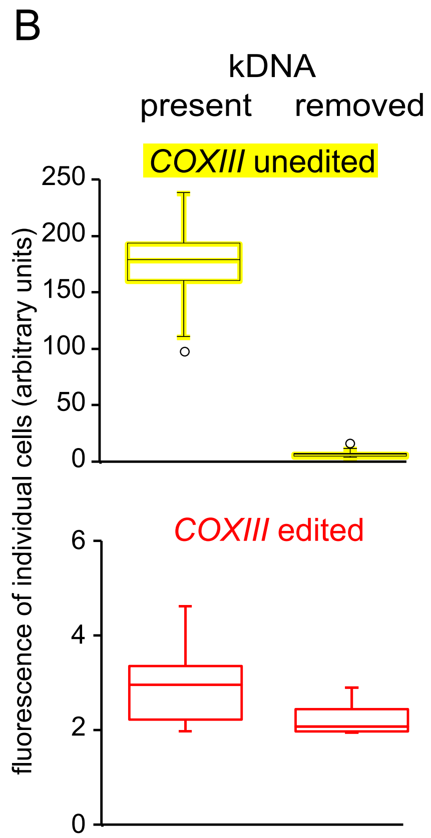


**Supplementary Figure S8: Specificity test for the *COXIII* probe set using akinetoplastic trypanosomes.**
BSF trypanosomes carrying a mutant of the ATPase gamma subunit (L262P) were treated with 10 nM ethidium bromide for three days to remove kDNA; untreated parasites served as control. Chemically fixed trypanosomes were probed via smFISH for edited and unedited *COXIII* mRNA as well as for *FUTSCH* (a cytoplasmic control mRNA).
**(A)** Projections (sum slices) of deconvolved Z-stacks (50 slices a 140 nm) are shown. Note the lack of kinetoplasts in the ethidium bromide samples (DNA stain) as well as the massive reduction in fluorescence for the unedited *COXIII* mRNA. In contrast, for the *FUTSCH* control mRNA, we only observed a mild reduction in the number of dots per cell (from 1.7±1 to 0.9±0.9; N>220), possibly caused by the minor effects 10 nM ethidium bromide have on nuclear DNA.
**(B)** The fluorescence of 10 individual cells (selected to each have two *FUTSCH* dots) was measured for edited and unedited *COXIII* and the data are presented as boxplot (waist is median; box is interquartile range (IQR); whiskers are ±1.5 IQR). The total fluorescence signal of cells probed for unedited *COXIII* is markedly reduced for akinetoplastic cells. Note that most cells do not have edited *COXIII* RNA, and therefore, no reduction with ethidium bromide is detectable.

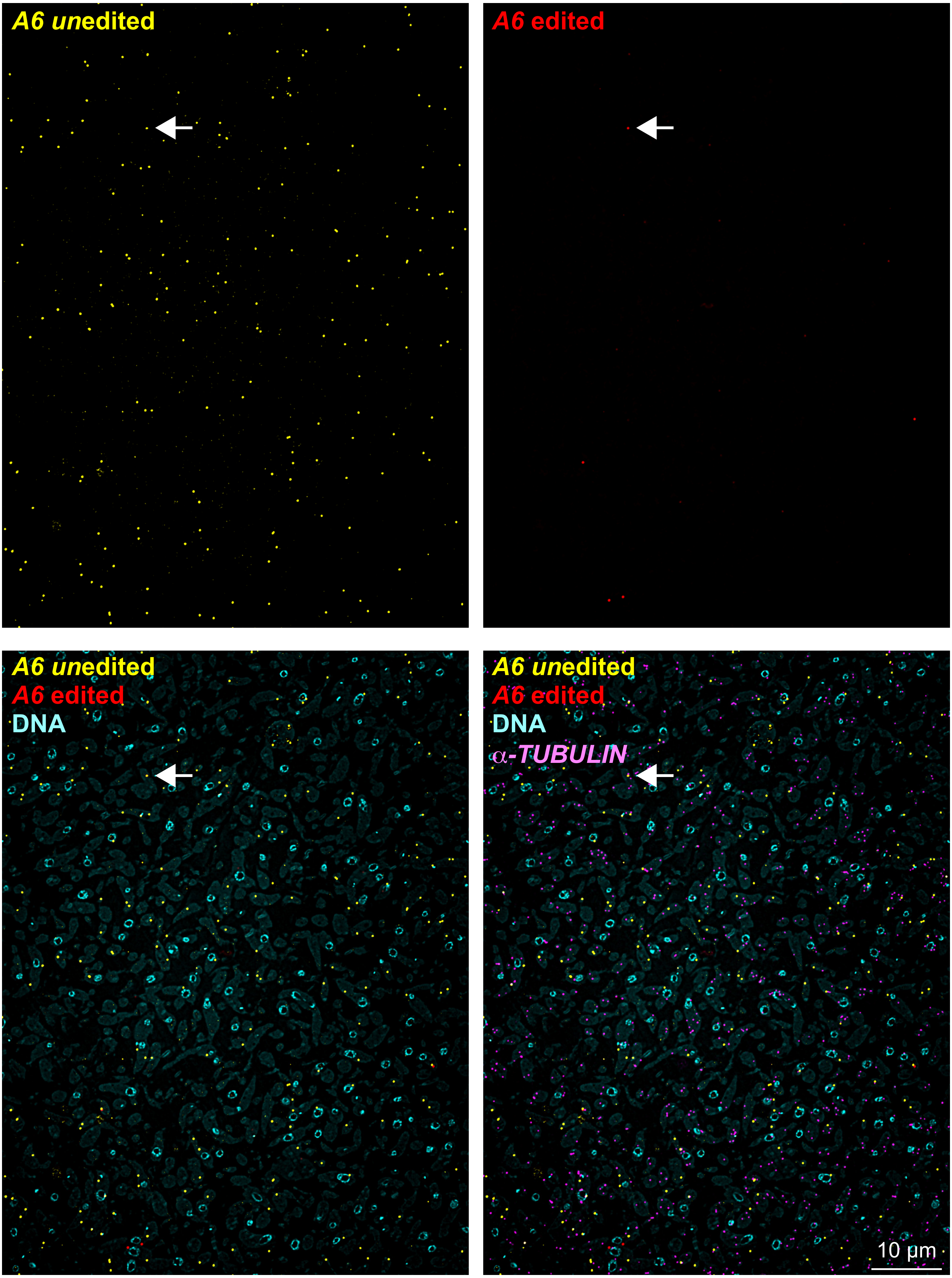


**Supplementary Figure S9: LR White probing for *A6***

LR White embedded samples of procyclic trypanosomes were probed as indicated. Projections of deconvolved Z-stacks are shown (sum slices, 5 slices a 140 nm). The white arrow points to a rare example of a double-stained spot (possibly a direct visualisation of RNA editing in progress).

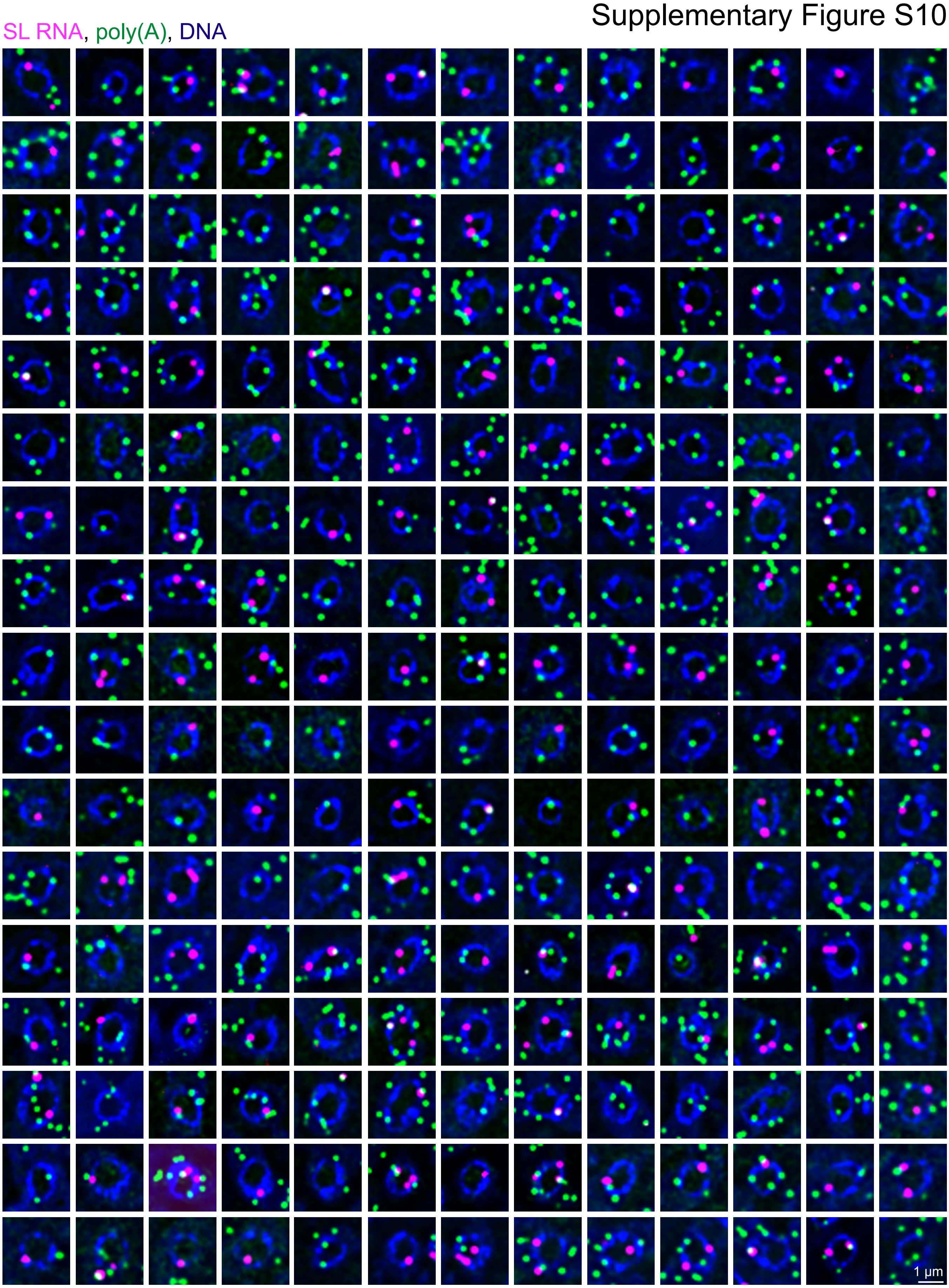


**Supplementary Figure S10: Quantification of nuclear-localised SL RNA and poly(A)**
smFISH was done on LR White embedded samples using a probe antisense to the SL RNA (pink) and to poly(A) (green). Nuclei of good shape (cut around the middle part rather than at the edge) were randomly chosen based on DAPI fluorescence only. All nuclei without any RNA in the nucleus were discarded, leaving the 221 nuclei shown here. From these, we quantified the number of nuclear-located SL RNAs (199) and poly(A) spots (355), which, corrected for the differences in the number of probe-pair recognition sites (2 for SL RNA and 0.94 for poly(A)) resulted in a 3.8 fold excess of nuclear poly(A) molecules in comparison to nuclear-localised SL RNA molecules.
